# Single-cell analyses reveal tissue-specific contextualization of HIF-1 activity

**DOI:** 10.1101/2024.07.02.601765

**Authors:** Ji Na Kong, D. Dipon Ghosh, Duanduan Ma, Ilona Kesisova, Vivek K. Dwivedi, Achilleus Savvidis, Steven R. Sando, Leigh Guerra, H. Robert Horvitz

## Abstract

Hypoxia (low oxygen) activates diverse cellular responses driven primarily by a single transcription factor, the hypoxia-inducible factor HIF-1. Here we identify at single-cell resolution HIF-1-dependent transcriptional responses in neuronal, intestinal, and muscle tissues of the nematode *Caenorhabditis elegans*. We report that while all three tissues share HIF-1-dependent modulation of pathways for central-carbon metabolism, individual tissues and specific cell subtypes exhibit broader transcriptional responses that differ extensively among cellular contexts. Using supervised machine learning to provide a deeper multivariate analysis, we observed that genotype-defined HIF-1 activation states were distinguishable within each tissue and that the highest-ranked predictive genes varied substantially among tissues. The differential-expression and machine-learning analyses together revealed a hierarchical contextualization of HIF-1-dependent transcriptomes: in response to HIF-1 activation, different tissues express distinct transcriptional programs, and cell subtypes within each tissue express further distinct programs. We validated representative tissue- and cell-type-specific responses *in vivo* and demonstrated that a gene that is HIF-1-responsive in a specific muscle cell subtype functions in that cell type to regulate HIF-1-driven hypersensitivity to the cholinergic agonist levamisole. Overall, our findings provide a systems-level view of HIF-1 activity and show that a broadly expressed transcription factor can generate both a shared core response and diverse context-dependent transcriptional programs.

## Main Text

Organismal responses to hypoxia (low oxygen) are critical for survival and adaptation. At the cellular level, the evolutionarily conserved prolyl hydroxylase EGLN (also referred to as EGL-9, PHD, or HIF-PH) and its target, members of HIF-1 (hypoxia-inducible) family of transcription factors, control responses to hypoxia^1–5^. For example, in normoxic conditions EGLN uses ambient oxygen to hydroxylate HIF-1ɑ, which marks HIF-1ɑ for proteasomal degradation. By contrast, in hypoxic conditions EGLN does not hydroxylate HIF-1ɑ, allowing stabilized HIF-1ɑ to enter the nucleus, regulate the expression of target genes that mediate the hypoxia response, and modulate myriad physiological and pathological processes, including glycolysis, angiogenesis, erythropoiesis, and tumorigenesis^5–8^. Although HIF-1ɑ is broadly expressed, hypoxia elicits distinct transcriptional and physiological responses in different tissues. Previous studies of *Caenorhabditis elegans* and vertebrate systems have identified cell type-specific HIF-1 targets and demonstrated that HIF-1 can influence physiology in both cell-autonomous and non-autonomous manners^5,9,10^. However, these studies have focused largely on individual genes or specific tissues; a systems-level view of how HIF-1-dependent transcriptional programs are implemented across multiple tissues within an intact animal has been lacking. For example, it remains unclear whether HIF-1 activation induces a largely shared transcriptional program that is incremented in relatively minor ways across tissues or instead engages distinct tissue-specific transcriptional responses. Here we report the HIF-1-dependent transcriptional responses of neuronal, intestinal, and muscle tissues of *C. elegans* based on multi-tissue single-cell RNA sequencing and Single-Cell Differential Expression Analysis (SCDE). We identify a conserved modulation of metabolic pathways shared across tissues coupled with highly divergent cell subtype-specific responses both within and among tissues. Supervised machine learning provided a deeper multivariate analysis and revealed that genotype-defined transcriptional states associated with HIF-1 activity can be distinguished based on an individual cell’s transcriptome and that the genes most predictive of HIF-1 activation state are conserved within tissues but vary substantially among tissues. Together, these findings provide a systems-level view of the tissue- and cell type-specific activity of the broadly expressed HIF-1 transcription factor in an intact animal.

## Generation of single-cell transcriptomic datasets for *C. elegans* neuronal, muscle and intestinal cells

We used *C. elegans* to investigate cell type-specific transcriptional responses driven by HIF-1 activity, because in *C. elegans*, in contrast with other animals, the *egl-9* and *hif-1* genes are not required for survival^4,11,12^. Furthermore, both the cellular anatomy and cell lineage of *C. elegans* have been completely described at the single-cell level^13,14^. We performed single-cell RNA sequencing (scRNA-Seq) analyses of three genotypes of *C. elegans* young (one-day) adults: wild-type, *egl-9* mutants, and *egl-9 hif-1* double mutants (Fig. 1a). *egl-9* mutant animals lack the prolyl hydroxylase that degrades HIF-1 in normoxic conditions; because HIF-1 is not degraded and therefore is constitutively active, these mutants function as if chronically exposed to hypoxia. By contrast, *egl-9 hif-1* double mutants lack HIF-1 function and can be used to identify which *egl-9*-dependent transcriptional changes require HIF-1 activation.

**Fig. 1.**
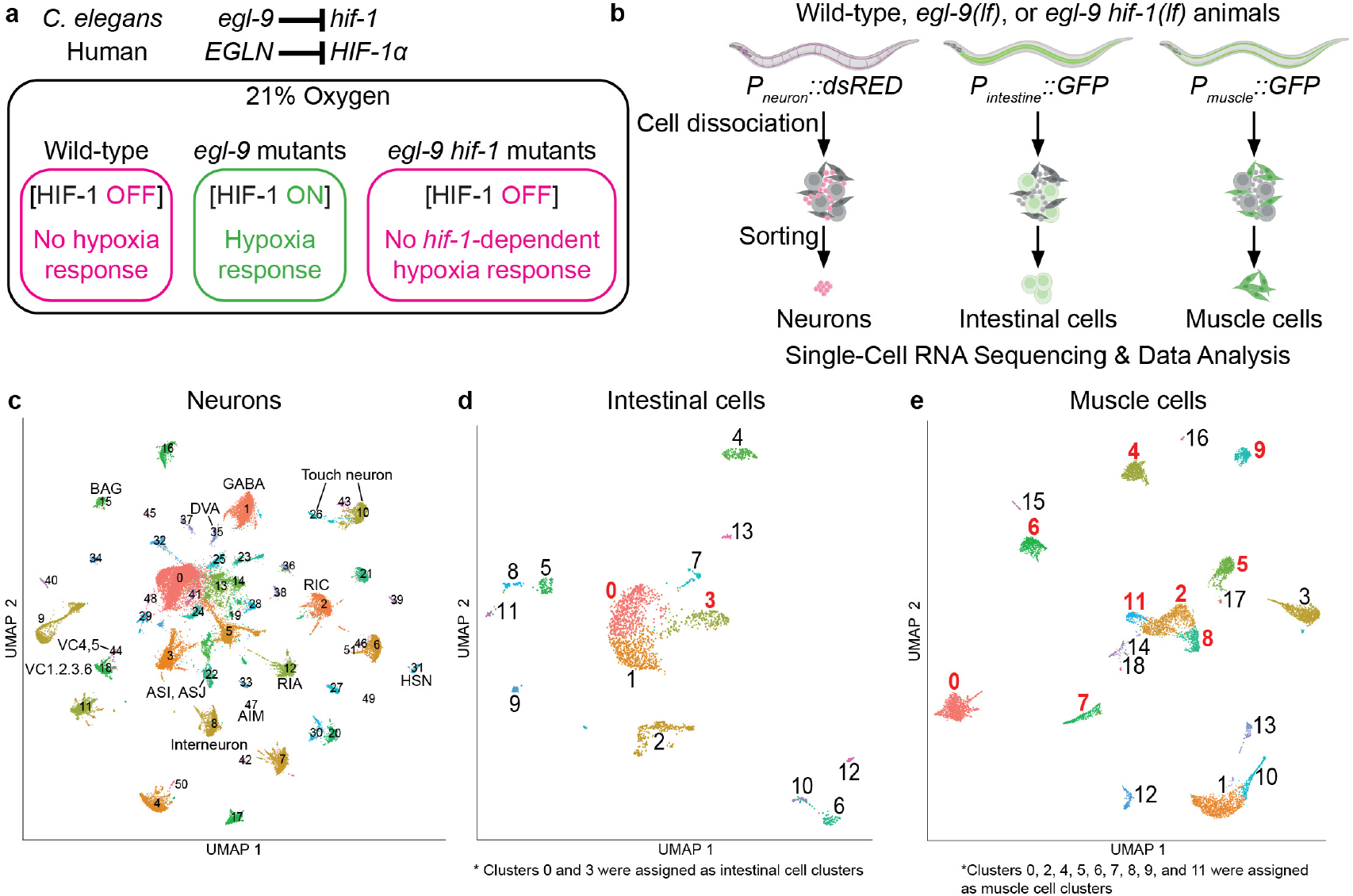
RNA-Seq analyses define HIF-1-dependent transcriptional responses across different tissues of *C. elegans* a, The evolutionarily conserved *egl-9-hif-1* pathway, in which the prolyl hydroxylase EGL-9 negatively regulates the hypoxia-inducible factor HIF-1. In *C. elegans*, loss-of-function mutations in *egl-9* cause constitutive HIF-1 activation and can serve as a proxy for chronic hypoxia-induced HIF-1 activation. Loss-of-function mutations in *hif-1* prevent the consequences of HIF-1 activation. **b,** Our experimental pipeline for scRNA-Seq analyses included the fluorescent labeling of tissues of interest, cell dissociation from whole animals, and sorting of target cells before subjecting them to transcriptomic analyses. We used the following promoters: neuronal *rgef-1*, intestinal *vha-6*, muscle *myo-3*^15–17^. **c-e** Combined UMAP projections of (**c**) neuronal, (**d**) intestinal, and (**e**) muscle cells. The identities of cells in clusters were assigned based on the expression of established cell-type-specific transcripts. Neuronal UMAP clusters were identified based on the expression of transcripts known to be specific to distinct neuron types^18^. Intestine UMAP clusters 0 and 3, highlighted in red, were assigned as intestinal cell clusters based on their expression of the known intestinal markers *vha-6* and *asp-1*^16,19^. Muscle UMAP clusters 0, 2, 4, 5, 6, 7, 8, 9, and 11, also highlighted in red, were assigned as muscle cell clusters based on their expression of the known muscle cell markers *myo-3* and *mup-2*^17^. See Extended Data Fig. 3 for more details.

We generated nine sets of mutant animals, one with each of the three genotypes (wild-type, *egl-9*, or *egl-9 hif-1*) expressing one of three fluorescent markers driven by a promoter specific to one of the three tissues of interest (neuronal *rgef-1*, intestinal *vha-6*, or muscle *myo-3*)^15–17^. To isolate specific cell types, we sorted dissociated target cells from the nine preparations of whole animals using fluorescence-activated cell sorting (FACS). We then performed nine rounds of scRNA-Seq to compare the gene-expression profiles of wild-type, *egl-9*, and *egl-9 hif-1* double-mutant animals for each of the three tissue types (Fig. 1b). We collected and analyzed a total of 35,913 neuronal cells, 2,509 intestine cells, and 9,085 muscle cells (Extended Data Fig. 1a). These numbers were sufficient to identify major intestinal and muscle cell populations and to capture diverse neuronal subtypes, although not all known neuronal classes were identified (Extended Data Fig. 1b-d). Detailed power calculations supporting these estimates are provided in the legend of Extended Data Fig.1.

We present a single Uniform Manifold Approximation and Projection (UMAP) projection of the three combined genotypes for a comprehensive overview of our scRNA-Seq datasets for neurons, intestinal cells, and muscle cells (Fig. 1c-e and Extended Data Fig. 2a-c). Using transcripts with characteristic and previously established expression patterns, we annotated the likely identities of numerous distinct clusters within our UMAP projections. Based on the expression of transcripts known to be specific to distinct neuron types^18^, we determined the identities of multiple neuronal clusters within our UMAP projections for neuronal tissue (Fig. 1c). Using a similar approach, we identified two intestinal cell clusters (clusters 0 and 3; Fig. 1d and Extended Data Fig. 3a,b) within our UMAP projections for intestinal tissue based on their expression of the established intestinal markers *vha-6* and *asp-1*^16,19^. Enrichment analysis of marker genes further suggests that intestinal cluster 7 corresponds to rectal gland and valve cells, and cluster 1 might represent a mixture of distal tip cells and intestine cells (data not shown). We identified 9 muscle-cell clusters (0, 2, 4, 5, 6, 7, 8, 9, and 11) (Fig. 1e and Extended Data Fig. 4c,d) within our UMAP projections for muscle cells based on expression of the established muscle-cell markers *myo-3* and *mup-2*^17^. In summary, our scRNA-Seq-based analysis enabled robust identification of neuronal, intestinal, and muscle cell types isolated from animals experiencing different levels of HIF-1 activity. These single-cell data establish a dataset for examining how HIF-1-dependent transcriptional responses differ among tissues and individual cell types in an intact animal.

## HIF-1-activated transcriptomes vary greatly among neuronal, intestinal, and muscle cells

To identify cell type-specific differences in gene expression among wild-type, *egl-9*, and *egl-9 hif-1* animals, we first used Single-Cell Differential Expression (SCDE) analyses^20^. We compared gene expression in cell-type clusters from *egl-9* mutants with expression in the corresponding clusters from wild-type animals to identify transcriptional changes caused by loss of *egl-9* function. Then, to identify those transcriptional changes specifically mediated by HIF-1, we determined which genes differentially regulated in *egl-9* mutants compared to wild-type animals were oppositely regulated in *egl-9 hif-1* mutant animals compared to *egl-9* mutants (Extended Data Table 1). We considered a transcriptional change as driven by HIF-1 activation if either (a) the gene was upregulated in *egl-9* mutants relative to wild-type animals and downregulated in *egl-9 hif-1* double mutants relative to *egl-9* mutants within the same cell cluster, or (b) the gene was downregulated in *egl-9* mutants relative to wild-type animals and upregulated in *egl-9 hif-1* double mutants relative to *egl-9* mutants within the same cell cluster (Fig. 2a).

**Fig. 2.**
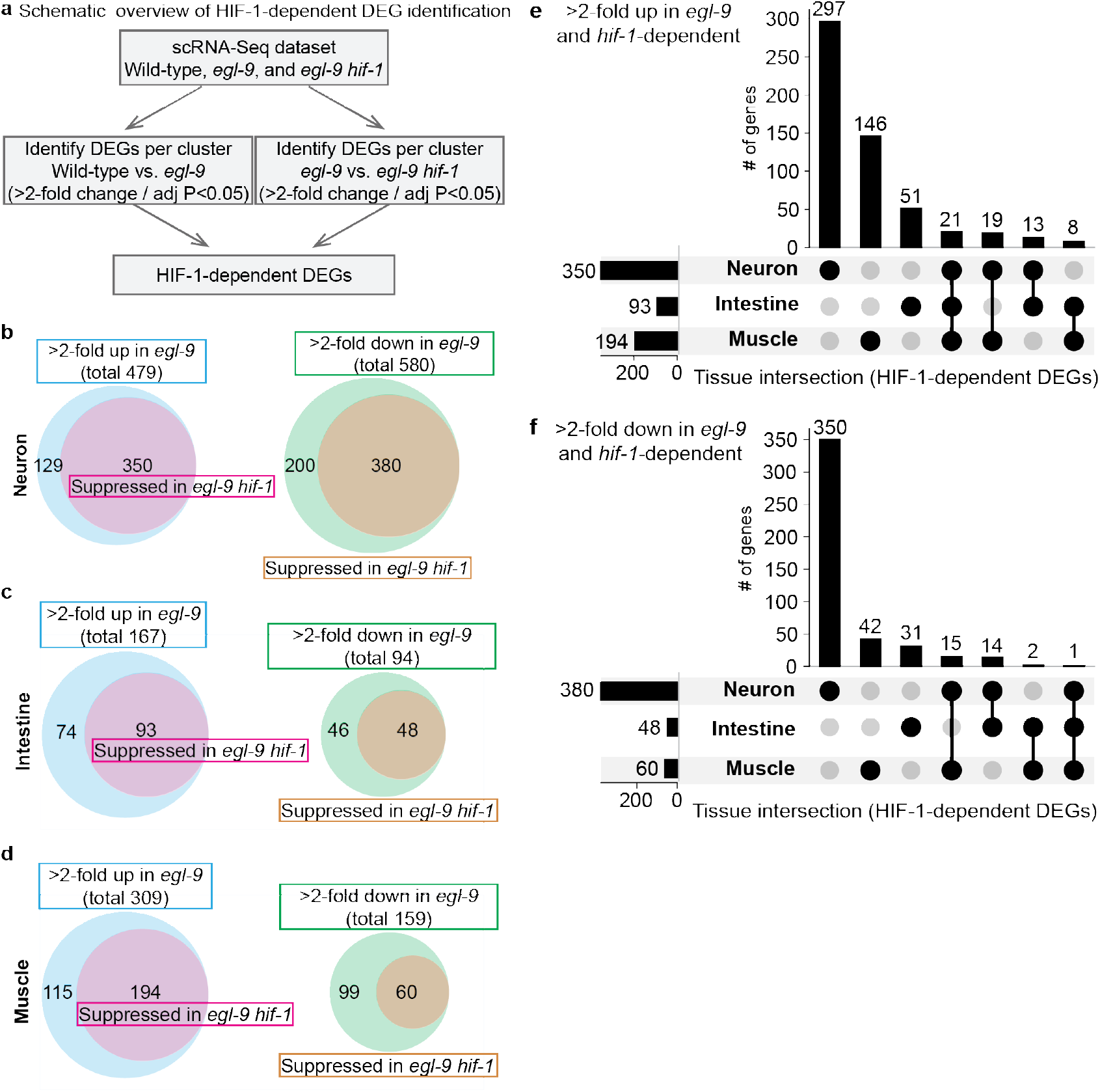
Overlapping and divergent HIF-1 activity among cell and tissue types reveals striking diversity in cellular hypoxia responses. **a**, Schematic overview of the strategy used to analyze differential gene expression among and within tissues. Differentially expressed genes (DEGs) were identified from two pairwise comparisons using scRNA-Seq data: wild-type vs. *egl-9* and *egl-9* vs. *egl-9 hif-1*, with adjusted *P* < 0.05. Genes significantly up- or downregulated in *egl-9* mutant cells and reciprocally down- or upregulated in *egl-9 hif-1* double mutant cells were defined as HIF-1-regulated. **b-d**, Venn diagrams illustrating the numbers of DEGs in neuronal (b), intestinal (c), and muscle (**d**) cells. Blue circles represent genes that were at least 2-fold upregulated in *egl-9* mutants compared to wild-type. Purple circles represent the subset of *egl-9* upregulated genes that were significantly downregulated in *egl-9 hif-1* mutants, indicating dependence on HIF-1 activity. Green circles represent genes that were at least 2-fold downregulated in *egl-9* mutants compared to wild-type. Beige circles represent the subset of *egl-9* downregulated genes that were significantly upregulated in *egl-9 hif-1* mutants, indicating dependence on HIF-1 activity. **e,** Intersection analysis of HIF-1-dependent genes upregulated in *egl-9* mutants across tissues. Bars indicate the number of genes shared among neuron, intestine, and muscle, with black circles identifying tissue combinations. Most HIF-1-dependent responses were tissue-specific, with limited overlap across all three tissues. **f**, Intersection analysis of HIF-1-dependent genes downregulated in *egl-9* mutants across tissues. As in (**e**), gene overlap across tissues was limited, indicating that both activation and repression programs downstream of HIF-1 are largely tissue-specific.

To assess the robustness of our scRNA-Seq analyses, we examined the expression of three well-established HIF-1 target genes – *pck-1*, *cysl-2*, and *nhr-57*^21–23^ – in neurons, intestine, and muscle cells of wild-type, *egl-9* mutant, and *egl-9 hif-1* double mutant animals. We observed significantly increased *pck-1*, *cysl-2*, and *nhr-57* expression in *egl-9* mutants compared both to wild-type and to *egl-9 hif-1* mutant animals in many cell clusters for all three tissues (Extended Data Fig. 4-7), consistent with previous bulk RNA-Seq studies^22–24^. Notably, different cells showed very different expression levels of *pck-1*, *cysl-2*, and *nhr-57* expression – a phenomenon not captured by whole-animal RNA-Seq approaches – indicating that our scRNA-Seq-based data offer new insights into the cell type-specific activity of even well-studied EGL-9/HIF-1-regulated genes and pathways.

We next identified genes expressed within neuronal, intestinal, and/or muscle tissues that exhibited significant differential regulation (>2-fold change / adjusted *p*-value < 0.05) in *egl-9* mutants compared to wild-type animals in a *hif-1-*dependent manner (Fig. 2b-d, Extended Data Table 1). These criteria thresholds were selected to identify genes exhibiting robust and biologically meaningful changes in expression and were established by previous transcriptomic studies^25–27^. We analyzed upregulated and downregulated genes separately and compared the resulting differentially expressed gene (DEG) lists across the three tissues (Fig. 2e, f, Extended Data Table 2). Strikingly, only 21 genes, including *pck-1*, *cysl-2*, and *nhr-57* (see below), exhibited differential HIF-1-mediated upregulation in at least one cluster of all three tissues. By contrast, most DEGs exhibited significant differential regulation in a tissue-specific manner. For example, 297 of the 350 genes upregulated in neurons were not upregulated in either intestinal or muscle cells (Fig. 2e). These DEG comparisons indicate that HIF-1-dependent transcriptional responses differ extensively among tissues.

## Genotype-defined transcriptional states associated with HIF-1 activity can be distinguished based on an individual cell’s transcriptome

Observing this divergence in HIF-1-responsive transcriptomes among tissues led us to analyze further the multigene patterns associated with HIF-1 activity. Differential-expression analyses identify HIF-1-dependent changes in the expression of individual genes within specific cell and tissue types. By contrast, a multivariate approach can examine patterns across many gene-expression features collectively within individual cells. We therefore asked whether the transcriptome of an individual cell contains sufficient information to distinguish genotype-defined transcriptional states associated with HIF-1 activity and, if so, which gene-expression features contribute to that distinction. To address these questions, we used supervised machine learning. We trained classifiers — algorithms that learned gene-expression patterns to distinguish cells with HIF-1 activity from cells without HIF-1 activity — on single-cell expression profiles separately for neuronal, intestinal and muscle cells (Fig. 3a). For each tissue, we performed two complementary binary genotype comparisons, wild-type versus *egl-9* and *egl-9* versus *egl-9 hif-1*, and evaluated nine classifiers and three input representations (normalized, square-root transformed, and rank-based) using five-fold cross-validation. Cells were divided randomly into five groups, known as “folds,” except that genotype proportions were preserved; in each of five iterations, four groups were used for training and the fifth for evaluation (see Methods). Hence, for the three tissues collectively, we performed a total of 810 training sessions (3 tissues X 2 genotypes X 9 classifiers X 3 input representations X 5 folds). To compare performance among tissues under a common modeling framework, we first used a neural-network classifier with the original Seurat-normalized expression representation from the differential-expression analysis. Mean cross-validated AUROC (Area Under the Receiver-Operating-Characteristic curve; 0.5 indicates chance-level discrimination, and 1 indicates perfect discrimination) for wild type versus *egl-9* was 0.999 for neurons (34,638 cells), 0.999 for intestinal cells (1,543 cells) and 0.990 for muscle cells (5,872 cells), with comparable values for *egl-9* versus *egl-9 hif-1* (0.999, 0.998 and 0.987, respectively) (Fig. 3b). High discriminatory performance was similarly observed across multiple linear, tree-based and neural-network classifiers for both genotype comparisons, whereas k-nearest-neighbor classifiers generally performed less well (Extended Data Fig. 8). In short, genotype-defined transcriptional states associated with HIF-1 activity could be distinguished from the transcriptomes of cells within each tissue. These results reveal that the transcriptomes of individual cells within each tissue contain sufficient information to distinguish genotype-defined transcriptional states associated with HIF-1 activity.

**Fig. 3.**
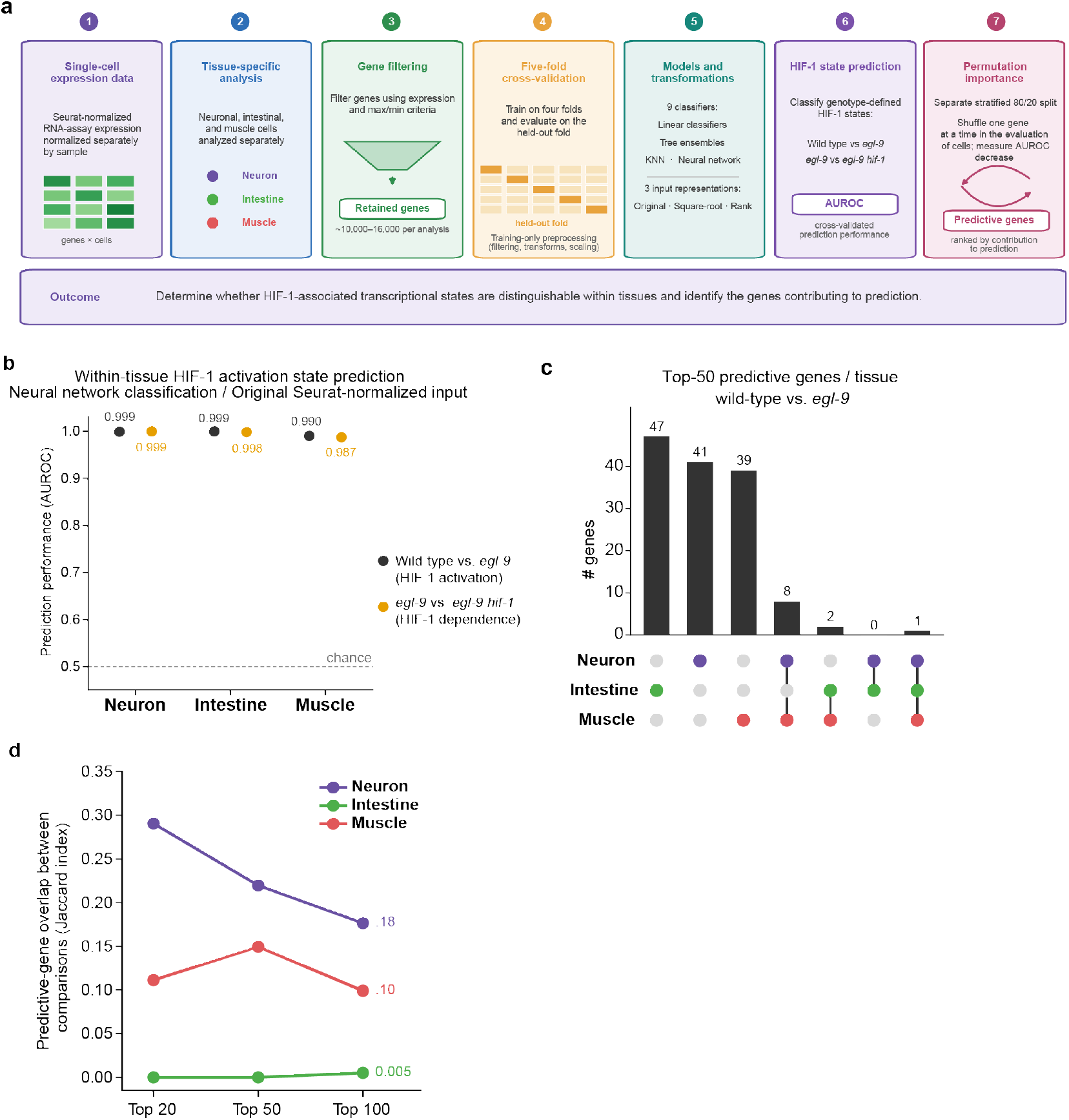
Machine-learning analysis identifies predictive transcriptional features of HIF-1 activation within tissues. **a**, Schematic of the within-tissue supervised machine-learning workflow. Seurat-normalized RNA-assay expression values were analyzed separately for neuronal, intestinal and muscle cells. Genes were filtered within training partitions, and nine classifiers combined with three input representations were evaluated using stratified five-fold cross-validation. Model performance was quantified by the area under the receiver-operating-characteristic curve (AUROC). Following model selection, permutation importance was calculated using a separate stratified 80/20 train/evaluation split to identify genes contributing to prediction. **b**, Within-tissue prediction of genotype-defined HIF-1 states using a common neural-network model with the original Seurat-normalized expression representation. AUROC values are shown for wild type versus *egl-9* and for *egl-9* versus *egl-9 hif-1* in neuronal (n = 34,638 cells), intestinal (n = 1,543 cells) and muscle cells (n = 5,872 cells). Points represent mean AUROC across five cross-validation folds; error bars indicate s.d. across the five folds and are smaller than the symbols. The dashed line indicates chance-level discrimination (AUROC = 0.5). **c**, Overlap among the top 50 predictive genes identified in neuronal, intestinal and muscle cells for the wild-type versus *egl-9* comparison. Predictive genes were ranked by mean permutation importance using the selected model–transformation configuration for each tissue (Extended Data Table 3). Permutation importance was defined as the decrease in AUROC following permutation of each gene in the evaluation subset. The UpSet plot shows the numbers of genes unique to or shared among tissue-specific top-50 predictive-gene sets. **d**, Overlap of predictive-gene sets between the two complementary genotype comparisons within each tissue. For neuronal, intestinal and muscle cells, the top 20, 50 and 100 genes ranked by permutation importance were compared between wild type versus *egl-9* and *egl-9* versus *egl-9 hif-1*. Overlap was quantified using the Jaccard index, defined as the number of shared genes divided by the number of genes in the union of the two sets. Exact gene-level comparisons for the top-50 sets are shown in Extended Data Fig. 9. For reference, Jaccard values expected under random selection were approximately 0.0006–0.0047 across the tissues and rank cutoffs examined.

## The genes most predictive of HIF-1 activation state differ among tissues, and many were not identified as differentially expressed in any individual cell type

What gene-expression features distinguish a cell’s genotype-defined HIF-1 state? We next asked how the gene-expression features supporting classification were distributed among tissues. Whereas differential-expression analysis identifies tissue-shared and tissue-specific responses using statistical tests that compare levels of individual genes, here we deployed the machine-learning method of permutation importance to assess the contribution of each gene to multivariate classification in the context of the remaining transcriptomic features. The expression values of each gene were randomly permuted across cells, disrupting that gene’s association with genotype while leaving all other features unchanged; genes for which permutation produced a larger reduction in AUROC were assigned greater predictive importance (Methods). For each tissue and genotype comparison, we ranked genes by permutation importance using the model– transformation configuration selected based on its high cross-validated AUROC (Methods; Extended Data Table 3). Among the top 50 predictive genes for the wild-type versus *egl-9* comparison, most were unique to a single tissue: 41 in neurons, 47 in intestine and 39 in muscle (Fig. 3c). Eight genes were shared between neurons and muscle, including the central-carbon-metabolism genes *pck-1*, *aldo-1* and *gpd-2*; two genes, *cysl-2* and *gst-19*, were shared between muscle and intestine; no genes were shared exclusively between neurons and intestine; and one gene, *F22B5.4*, a previously identified HIF-1-dependent hypoxia-responsive gene^28^, was among the top 50 in all three tissues. *F22B5.4* was also among the 21 genes we identified to be upregulated in a HIF-1-dependent manner in all three tissues by differential-expression analysis (Fig. 2e), providing an example of concordance between the two analytical approaches. The limited overlap among the three tissue top-50 sets indicates a tissue contextualization of the features contributing to prediction, although exact overlap estimates should be interpreted cautiously because the selected model–transformation configurations differed among tissues and permutation importance can be affected by correlations among gene-expression features.

We next asked whether predictive features were conserved across the two complementary genotype comparisons within each tissue. Across the top 20, 50 and 100 predictive genes, overlap (Jaccard index; 0, no overlap; 1, complete overlap) was consistently greatest in neurons (0.18–0.29), intermediate in muscle (0.10–0.15) and minimal in intestine (0–0.005) (Fig. 3d). At the top-50 cutoff, 18 genes were shared between the two neuronal comparisons and 13 between the two muscle comparisons, whereas no top-50 genes were shared between the intestinal comparisons (Extended Data Fig. 9). Predictive genes identified in both genotype comparisons within each tissue included *F22B5.4*, *pck-1* and *glb-1* in both neurons and muscle; *mdh-1* and *enol-1* in neurons; and *gpd-2*, *gpd-3* and *pcca-1* in muscle. In neurons, this overlap was observed even though different model–transformation configurations were selected for the two comparisons. The intestinal comparisons showed minimal overlap across all three cutoffs; extensive ties and variability in permutation-importance values around the rank cutoffs make the exact membership of the intestinal top-gene sets particularly sensitive to small ranking differences. Exact overlap estimates should be interpreted cautiously because the selected model-transformation configurations differed among tissues and genotype comparisons, and permutation importance can be affected by correlations among gene-expression features. In short, we identified subsets of predictive genes that were shared between the wild-type versus *egl-9* and *egl-9* versus *egl-9 hif-1* comparisons in neurons and muscle; this overlap was minimal in intestine.

We then assessed the relationship between the predictive genes identified by machine learning and the genes identified by the cluster-specific differential-expression data. We compared the top 100 predictive genes with the genes identified by cluster-wise differential-expression analysis. Among the 100 most-predictive genes in the wild-type versus *egl-9* comparison, some—27% in neurons, 56% in muscle and 82% in intestine—were not identified as differentially expressed in any individual cell cluster (Extended Data Table 3). In other words, the predictive-gene rankings overlapped only in part with the cluster-wise DEG lists, and the extent of overlap differed among tissues. These findings reveal that the tissue contextualization of HIF-1 activation state identified by multivariate machine-learning analysis extends well beyond the genes that meet cluster-wise differential-expression thresholds. These results establish that supervised learning provides an informative complementary approach to the study of transcriptional variation.

Together these analyses establish that genotype-defined transcriptional states associated with HIF-1 activity are distinguishable within each tissue and that the highest-ranked features predictive of genotype differ substantially among tissues. Only some of the predictive genes shared across tissues or between the two complementary genotype comparisons were also identified in the shared HIF-1-dependent response as defined by differential-expression analysis (Fig. 2e). More broadly, these results illustrate how multivariate supervised learning can complement differential-expression analyses by capturing collective patterns across multiple gene-expression features within individual cells, thereby providing an integrated view of how HIF-1 activity is represented in different tissue contexts.

## *In vivo* validation of tissue-specific HIF-1-responsive genes

To test the *in vivo* expression patterns of genes identified as tissue-specific and HIF-1-dependent by our scRNA-Seq analyses, we examined representative genes from the neuronal, intestinal, and muscle datasets using single-molecule Hybridization Chain Reaction fluorescence *in situ* hybridization (HCR smFISH) and transcriptional reporters. Candidate genes were chosen based on strong tissue-restricted differential expression patterns in the scRNA-Seq dataset. We excluded broadly expressed HIF-1 targets such as *cysl-2* and *pck-1*. Within each tissue, we selected genes representing both broadly distributed and more spatially restricted cluster-specific expression patterns.

We examined 12 genes using HCR smFISH: four neuronal (*scl-19*, *T10B9.9*, *gcy-33*, and *gcy-29*), three intestinal (*ftn-1, mtl-1*, and *oac-54*) and five muscle (*mmcm-1*, *pcca-1*, *pccb-1*, *cpg-3*, and *mct-6*). Nine of the 12 genes showed tissue-enriched expression that was increased in *egl-9* mutants and reduced in *egl-9 hif-1* double mutants, consistent with HIF-1-dependent regulation. *T10B9.9* and *gcy-33* showed neuron-enriched HIF-1-dependent induction in subsets of neurons, although the specific neuronal identities were not determined (Extended Data Fig. 10a,b). *scl-19* also showed neuron-enriched induction by HIF-1 and was broadly distributed across neuronal subtypes (data not shown). *gcy-29* did not yield detectable signal. *ftn-*1, *mtl-1*, and *oac-54* each showed intestine-enriched induction by HIF-1 (Extended Data Fig. 10c-e). *mmcm-1*, *pcca-1*, and *pccb-1* showed muscle-enriched induction by HIF-1 (Extended Data Fig. 10f-h). By contrast, *cpg-3* was induced by HIF-1 predominantly in the germline rather than muscle, and *mct-6* expression could not be confidently localized (data not shown).

To test these responses using an independent method, we generated transcriptional reporters and obtained interpretable expression patterns for eight genes that showed HIF-1-dependent induction: *aqp-7* in neurons; *ftn-1*, *mtl-1*, and *cysl-3* in intestine; and *mmcm-1*, *pcca-1*, *gpd-3*, and *pccb-1* in muscle (Extended Data Figs. 11–13). HCR smFISH and transcriptional reporters were used as independent approaches to examine overlapping but non-identical sets of genes, and concordance between the methods could be assessed for five genes—*ftn-1*, *mtl-1*, *mmcm-1*, *pcca-1*, and *pccb-1*. All five showed HIF-1-dependent induction by both HCR smFISH and analyses of transcriptional reporters. Across both methods, we examined 15 genes and obtained experimental support for the predicted tissue-enriched HIF-1-dependent induction of 12. Of the remaining three genes, *gcy-29* did not yield a detectable signal, *mct-6* expression could not be confidently localized, and *cpg-3* was detected predominantly in the germline rather than in muscle.

In several instances, the scRNA-Seq dataset also detected lower-level differential expression in additional tissues — for example, *mtl-1* and *pcca-1* in neurons, *cysl-3* in muscle, and *gpd-3* across neurons, intestine, and muscle (Extended Data Table 2). The imaging analyses primarily detected expression in those tissues exhibiting the strongest expression and did not consistently reveal the weaker changes detected by scRNA-Seq in additional tissues. We therefore interpret these experiments as tests of the predominant anatomical sites of HIF-1-dependent induction rather than as comprehensive measurements of lower-level expression across all tissues.

Data for 11 of these 12 genes are shown in Extended Data Figs. 10–13; *scl-19*, which also showed neuron-enriched HIF-1-dependent induction, was broadly expressed and for that reason is not represented in the figures. Of the 11 genes with clear tissue-specificities, nine had previously been identified as HIF-1-regulated from bulk RNA-Seq datasets^22,24,29^, and two genes, *T10B9.9* and *aqp-7*, were newly identified through our single-cell analyses. Neither *T10B9.9* nor *aqp-7* was identified as differentially expressed in any of the three independent bulk RNA-Seq datasets we examined, yet both showed clear neuron-enriched HIF-1-dependent induction *in vivo* (Extended Data Figs. 10, 11). For the genes previously identified by bulk RNA-Seq, our imaging analyses additionally defined their predominant anatomical sites of regulation—for example, *gcy-33* in neurons, *ftn-1* in intestine, and *mmcm-1* in muscle.

In short, these findings provide experimental validation of the tissue- and cell-type-specific HIF-1-dependent transcriptional responses identified by the scRNA-Seq dataset.

## Nearly all of the shared HIF-1-responsive genes but fewer than half of the tissue-specific genes show evidence consistent with direct HIF-1 regulation

We next asked whether the HIF-1-dependent DEGs identified in our atlas show evidence consistent with direct HIF-1 regulation. To assess potential direct HIF-1 binding, we compared our DEGs with a genome-wide HIF-1 ChIP-Seq dataset^22^ available through ENCODE^30^ that used an anti-GFP antibody to map HIF-1 occupancy in whole animals expressing HIF-1::GFP. A DEG was scored as having evidence for HIF-1 binding if it was associated with a HIF-1 ChIP-Seq peak. In parallel, we searched the proximal promoter regions (≤1 kb upstream of the transcription start site) of each DEG for the canonical hypoxia response element (HRE) sequence, RCGTG. Because this short consensus sequence occurs frequently by chance, and because functional HREs can also reside within introns, 3′ untranslated regions (UTRs), or other regulatory elements^22,31^, we treated ChIP-Seq occupancy as the primary evidence of HIF-1 binding and a promoter-proximal HRE as supporting evidence for DEGs with ChIP-Seq occupancy.

Among the 21 genes upregulated across all three tissues, 20 showed HIF-1 ChIP-Seq binding and contained promoter-proximal HREs (Extended Data Fig. 14a,b, Extended Data Table 2). The only exception, *swm-1,* was previously identified as upregulated in bulk RNA-Seq datasets^24^ and contains a canonical HRE but lacks clear ChIP-Seq evidence. Thus, nearly all genes induced across all three tissues have strong evidence consistent with direct HIF-1 regulation.

We next examined genes upregulated specifically in one tissue: 297 neuronal, 51 intestinal, and 146 muscle genes (Fig.2e). A HIF-1 ChIP-Seq peak was present at 22% of neuron-specific, 31% of intestine-specific, and 41% of muscle-specific upregulated genes (Extended Data Fig. 14d, Extended Data Table 2). Promoter HREs were more common, occurring in 60–73% of these genes, but were also common among tissue-specific downregulated genes, which are likely to be indirectly regulated by HIF-1 (see below). Thus, we did not consider HRE presence alone to be sufficient evidence for direct regulation. These results indicate that about 20-40% of the tissue-specific upregulated genes are likely to be regulated directly by HIF-1. Genes lacking detectable ChIP-Seq binding might be regulated indirectly or through tissue-restricted HIF-1 binding not detected from the available whole-animal dataset.

Only one gene, *ahcy-1*, was downregulated in a HIF-1-dependent manner across all three tissues. *ahcy-1* contained a promoter HRE but lacked a HIF-1 ChIP-Seq peak (Extended Data Fig. 14c, Extended Data Table 2). We applied the same analysis to genes downregulated specifically in neurons, intestine, or muscle: 350 neuronal, 31 intestinal, and 42 muscle genes.

HIF-1 ChIP-Seq occupancy was less frequent among downregulated than upregulated genes in all three tissues, occurring in 7%, 23%, and 10% of neuron-, intestine-, and muscle-specific downregulated genes, respectively (Extended Data Fig. 14d, Extended Data Table 2). By contrast, promoter HRE frequencies were similar among tissue-specific upregulated and downregulated genes. Overall, HIF-1 ChIP-Seq binding was associated with 140 of 494 tissue-specific upregulated genes (28%) but only 34 of 423 tissue-specific downregulated genes (8%). The relatively low frequency of HIF-1 binding among downregulated genes is consistent with the downregulation of HIF-1-dependent DEGs occurring predominantly through indirect downstream mechanisms.

Together, these analyses indicate distinct patterns of HIF-1-binding among HIF-1-responsive genes. Genes upregulated across all three tissues are strongly associated with direct HIF-1 binding, whereas tissue-specific upregulated genes include both genes with and without detectable binding. HIF-1 binding was comparatively uncommon among downregulated genes, suggesting that much of HIF-1-dependent downregulation is indirect. Because the ChIP-Seq data were generated from whole animals expressing a HIF-1::GFP transgene, an associated peak indicates HIF-1-binding potential at a locus but does not establish occupancy in the specific tissue or physiological context in which the gene was differentially expressed in our experiments.

## Genes induced by HIF-1 in all three tissues are enriched for central carbon metabolism

Previous whole-animal bulk RNA-Seq studies of *C. elegans* have established that HIF-1 modulates various metabolic (e.g., glucose-related) and immune-related processes^22,24^. Having identified HIF-1-dependent DEGs at single-cell resolution across neuronal, intestinal, and muscle tissues, we next asked which biological processes are preferentially engaged in each tissue. We first examined the set of 21 genes upregulated in a HIF-1-dependent manner in all three tissues (Fig. 2e). WormCat^32^ and Gene Ontology (GO) enrichment analyses of this shared set both identified enrichment for metabolic categories (Fig. 4b; Extended Data Tables 4 and 5). Ten of the 21 shared genes were assigned to metabolism (Bonferroni FDR = 9.0 × 10⁻⁶), and the most significant enrichment (WormCat category “glycolysis”; FDR = 1.6 × 10⁻¹¹) corresponded to six genes that encode central-carbon-metabolism enzymes: the glycolytic enzymes *aldo-1*, *enol-1*, *gpd-2*, and *gpd-3*, the gluconeogenic enzyme *pck-1*, and the TCA-cycle enzyme *mdh-1*. Because these genes are upregulated in all three tissues, the shared response reflects a common central-carbon-metabolism module rather than distinct tissue-specific metabolic programs and is consistent with the metabolic regulation described in whole-animal bulk studies^22,28,33^. Genes shared between only two of the three tissues showed enrichment limited to specific Cat2 modules, including lipid metabolism (neuron–intestine), extracellular matrix components (neuron–muscle), and nucleotide/stress-associated pathways (intestine–muscle), indicating that inter-tissue overlap is pathway-specific rather than broadly homogeneous (Extended Data Fig. 15).

**Fig. 4.**
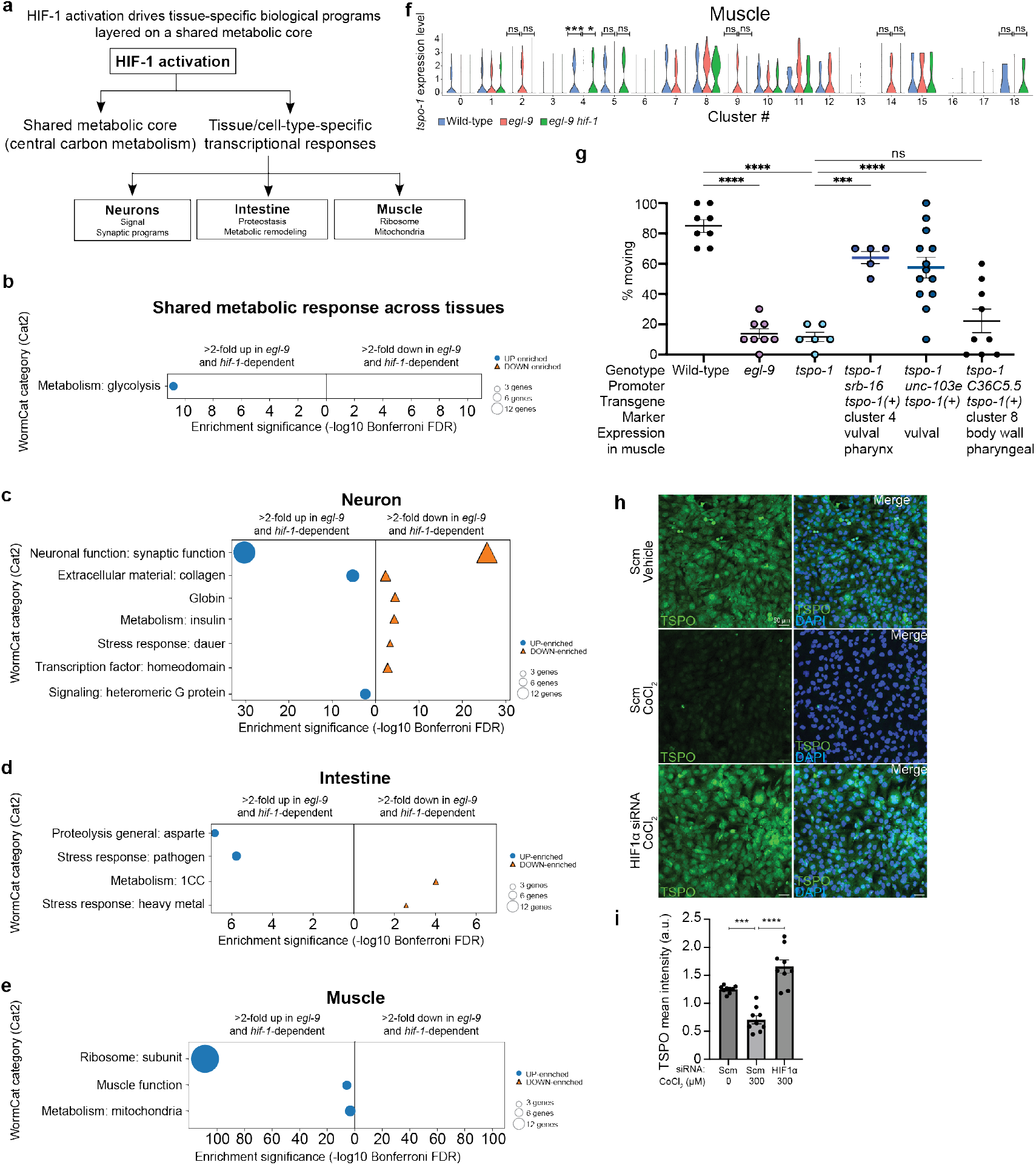
Tissue- and cell type-specific HIF-1 programs drive distinct biological responses. a, Schematic showing that HIF-1 activation is associated with both shared and tissue-specific transcriptional responses. A shared metabolic response, comprising central carbon metabolism genes, is observed among neuronal, intestinal, and muscle tissues, whereas additional tissue-specific programs differ among both tissue and cell types. **b,** WormCat enrichment analysis of HIF-1-dependent genes shared among all three tissues (n = 21). The WormCat “glycolysis” category comprises central-carbon-metabolism enzymes (glycolytic *aldo-1*, *enol-1*, *gpd-2*, *gpd-3*; gluconeogenic *pck-1*; TCA *mdh-1*). Analyses were performed using genes upregulated in *egl-9* mutants and reciprocally downregulated in *egl-9 hif-1* double mutants. **c–e,** Tissue-specific WormCat enrichment analyses of HIF-1-up- or downregulated genes in neurons (c), intestine (d), and muscle (e). Enriched functional categories differ among tissues, indicating that HIF-1 activation engages distinct biological programs in different cellular contexts. Neuronal responses are enriched for synaptic and signaling-related functions, intestinal responses for proteolysis and pathogen-stress responses, and muscle responses for ribosome biogenesis and mitochondrial metabolism. For all enrichment analyses (b-e), dot size reflects the number of genes in each category and color and shape indicate direction of regulation. Enrichment significance is shown as –log_10_ Bonferroni-adjusted false discovery rate. **f,** Violin plots depicting *tspo-1* expression levels in all muscle-cell clusters in wild-type (blue), *egl-9* mutant (red), and *egl-9 hif-1* double-mutant (green) animals. Among muscle clusters, statistically significant HIF-1-dependent changes in *tspo-1* expression were observed specifically in muscle cluster 4, identified as vulval muscle cells (see Extended Data Fig. 23e). Although several additional clusters exhibited detectable *tspo-1* expression, these clusters did not show statistically significant HIF-1-dependent regulation. These analyses indicate that HIF-1-dependent modulation of *tspo-1* occurs in a cell-type-restricted manner within muscle tissue. **g,** *egl-9* and *tspo-1* null mutants are hypersensitive to 2 mM levamisole. Levamisole is a potent nicotinic acetylcholine receptor agonist that stimulates *C. elegans* muscle contraction and can lead to immobility. Each data point represents one independent assay of ∼10 adult animals. Approximately 10 adult worms were exposed to the specified concentrations of levamisole, and the number of animals moving was scored after 10 min of exposure. Quantification of sensitivity to 2 mM levamisole of *tspo-1* null mutants with the wild-type *tspo-1*(+) gene expressed specifically in cluster 4 cells using the *srb-16* promoter, vulval muscle cells using the *unc-103e* promoter, and cluster 8 cells (which do not include vulval muscle cells) using the *C36C5.5* promoter. Expression of *tspo-1* in muscle cluster 4 cells or vulval muscle cells rescued the levamisole hypersensitivity of *tspo-1* null mutants. By contrast, expression of *tspo-1* in muscle cluster 8 cells did not rescue the levamisole hypersensitivity of *tspo-1* null mutants. Bars indicate mean ± s.e.m. ns, not significant; \**P* < 0.05, \*\**P* < 0.01, \*\*\**P* < 0.001, \*\*\*\**P* < 0.0001. **h,i** Immunocytochemistry (h) and corresponding quantification (i) of TSPO levels in AC16 cardiomyocytes treated with cobalt chloride (CoCl_2_) in the presence or absence of HIF-1α siRNA. TSPO is shown in green and DAPI in blue; scale bar, 50 µm. Cobalt chloride treatment reduced TSPO levels, and this reduction was suppressed by HIF-1α knockdown. Mean fluorescence intensity was quantified per image and normalized to DAPI; each data point represents one image. Bars indicate mean ± s.e.m. \*\*\**P* < 0.001, \*\*\*\**P* < 0.0001.

The shared modules together accounted for only a minority of the HIF-1-dependent DEGs in each tissue. We therefore next examined the genes regulated in a tissue-restricted manner to determine what distinguishes the HIF-1 response among neurons, intestine, and muscle.

## Neuron-specific HIF-1 activation modulates synaptic and G-protein signaling programs

WormCat analysis of neuron-specific HIF-1-dependent DEGs revealed strong enrichment for neuronal-function categories, particularly synaptic function, which was driven largely by neuropeptide signaling. Additional enriched categories included extracellular matrix/collagen components and heteromeric G-protein signaling, together with downregulated globin, insulin-like peptide, dauer-larva pathway, and homeodomain transcription-factor categories (Fig. 4c; Extended Data Table 4). Both upregulated and downregulated gene sets contributed to the synaptic and extracellular-matrix enrichments, indicating bidirectional modulation of neuronal signaling and structural programs following HIF-1 activation. GO enrichment analysis yielded concordant results, with particularly strong enrichment for neuropeptide signaling and additional enrichment for G-protein-coupled receptor signaling and extracellular and cuticle structural terms (Extended Data Fig. 16; Extended Data Table 5). Together, these analyses indicate that neuronal HIF-1 activation is associated with coordinated, bidirectional modulation of neuropeptide signaling and extracellular-matrix genes, together with changes in G-protein signaling.

## Intestine-specific HIF-1 activation modulates metabolic and stress-response programs

Consistent with the function of intestinal tissue as a protective barrier and metabolic hub in the worm^34,35^, WormCat enrichment analysis of intestine-specific HIF-1-dependent DEGs revealed distinct enrichment patterns between upregulated and downregulated genes.

Upregulated genes were enriched for aspartyl proteases and pathogen-response genes, whereas downregulated genes were strongly enriched for metabolic categories, including one-carbon and transsulfuration metabolism; heavy-metal-response genes were also enriched among the downregulated genes (Fig. 4d; Extended Data Table 4). The downregulated metabolic set included genes associated with fatty-acid oxidation and gluconeogenesis. GO analysis of the up- and downregulated combined intestine-specific DEG set yielded complementary results, identifying enrichment for fatty-acid oxidation, glucose and pyruvate metabolism, amino-acid metabolism, innate immune and defense responses, and aspartic-type peptidase activity (Extended Data Fig. 16; Extended Data Table 5). Together, these analyses indicate that intestinal HIF-1 activation alters the expression of genes associated with proteolysis, pathogen responses, and multiple metabolic processes, with proteolytic and pathogen-response categories enriched among upregulated genes and metabolic categories enriched among downregulated genes.

## Muscle-specific HIF-1 activation modulates translational and mitochondrial programs

WormCat analysis of muscle-specific HIF-1-dependent DEGs revealed particularly strong enrichment for ribosome-related categories among upregulated genes, together with enrichment for muscle function and mitochondrial metabolism categories (Fig. 4e; Extended Data Table 4). The enriched muscle-function genes included components associated with muscle structure and contraction, whereas the mitochondrial category included genes associated with oxidative metabolism. GO enrichment analysis of the up- and downregulated combined muscle-specific DEG set yielded complementary results, identifying enrichment for cytoplasmic translation and ribosome-related categories, mitochondrial electron transport and respiration, and muscle structural components including sarcomeres and myofibrils (Extended Data Fig. 16; Extended Data Table 5). Together, these analyses indicate that muscle HIF-1 activation is associated with transcriptional changes in ribosomal, mitochondrial-metabolic, and contractile gene programs.

## Tissue-specific HIF-1 responses include strongly induced genes

Having identified distinct tissue-specific programs, we next asked whether tissue-specific responses were limited to genes with relatively weak induction. We compared the induction magnitudes of genes upregulated specifically in each tissue with those of genes upregulated across all three tissues (Extended Data Fig. 17; Extended Data Table 4). Within each tissue, genes upregulated across all three tissues showed higher median *egl-9* versus wild-type log2-fold changes than tissue-specific upregulated genes (median log2-fold changes of 2.7, 2.7, and 2.5 for the 21 shared genes versus 1.6, 1.9, and 1.6 for tissue-specific genes in neurons, intestine, and muscle, respectively; Extended Data Fig. 17). Shared genes were also enriched among the 20 most strongly induced genes; they constituted 25%, 45%, and 55% of the top 20 genes in neurons, intestine and muscle, respectively, compared with 6%, 23%, and 11% of all HIF-1-dependent upregulated genes in those tissues, corresponding to approximately two-to fivefold enrichment. Nevertheless, tissue-specific genes were represented among the 20 most strongly induced genes in all three tissues—15 in neurons, 8 in the intestine, and 5 in muscle—and included *T10B9.9*, the gene most strongly induced in neurons. Thus, although genes shared across all three tissues were more strongly induced on average, tissue-specific HIF-1 responses were not restricted to weakly induced genes. Together, these findings indicate that a strongly induced shared transcriptional core is accompanied by high-magnitude tissue-specific responses, highlighting the diversity of cellular responses to hypoxia within a single organism.

## Cell subtypes within tissues exhibit divergent HIF-1-dependent transcriptional responses

We next asked whether HIF-1-dependent transcriptional responses vary among cell subtypes within individual tissues. GO enrichment analyses of HIF-1-dependent DEGs from selected neuronal, intestinal, and muscle clusters revealed both shared and distinct functional enrichment patterns (Extended Data Figs. 18-20; Extended Data Table 5). Because the numbers of HIF-1-dependent DEGs varied substantially among clusters and could influence the number of significant GO terms detected, we compared the identities of the enriched functional categories rather than the numbers of significant terms.

For example, within the nervous system, BAG neurons (cluster 15), which detect decreasing concentrations of oxygen^36^, and touch receptor neurons (clusters 10 and 26) shared enrichment for pyruvate metabolism (Extended Data Figs. 18; Extended Data Table 5), whereas neuropeptide signaling was additionally enriched in BAG neurons but not in the two touch-receptor clusters.

Two intestinal cell clusters (cluster 0 and cluster 3, the latter likely representing posterior intestinal cells based on marker expression; Extended Data Fig. 21) exhibited substantial overlap in metabolic and stress-response enrichment. Nevertheless, whereas cluster 0 showed strong enrichment for immune-related processes, cluster 3 displayed enrichment for sulfur amino acid metabolic and biosynthetic processes (Extended Data Fig. 19; Extended Data Table 5). These findings suggest that, although several HIF-1-responsive metabolic and stress-response processes are shared, posterior intestinal cells might engage regionally specialized metabolic programs. Such intra-tissue heterogeneity could reflect differences in the physiological roles of distinct intestinal regions.

Analyses of clusters corresponding to vulval muscle (cluster 4; Extended Data Fig. 22a) and anterior body-wall muscle (cluster 8; Extended Data Fig. 22b) revealed enrichment for different functional categories. Cluster 4 was enriched for translational and biosynthetic processes, including cytoplasmic translation, ribosomal small subunit biogenesis, and gene expression, as well as muscle contraction–related terms. By contrast, cluster 8 was enriched for organic-acid and carbohydrate metabolism, including oxaloacetate, propionate, and glucose metabolism; mitochondrial-matrix components were also enriched (Extended Data Fig. 20; Extended Data Table 5). Thus, the HIF-1-dependent DEG sets in anatomically distinct muscle subtypes were associated with different functional categories. The functional significance of the translational and biosynthetic enrichment in vulval muscle remains to be determined.

These analyses indicate that HIF-1-dependent transcriptional responses differ not only among tissues but also among selected cell subtypes within individual tissues.

## An evolutionarily conserved HIF-1-responsive pathway acting within a specific muscle cell subtype can modulate *C. elegans* physiology

To determine if tissue subtype-specific HIF-1-responsive transcriptional programs identified by our atlas function physiologically, we sought HIF-1 effectors that influence muscle function. We observed that *egl-9* mutants are hypersensitive to the cholinergic agonist levamisole, which induces sustained muscle contraction and immobility^37–39^ (Extended Data Fig. 23a,b), and that this hypersensitivity was suppressed in *egl-9 hif-1* double mutants (Extended Data Fig. 23a), indicating that the phenotype is HIF-1 dependent. These findings indicate that HIF-1 activation alters levamisole sensitivity. We then tested 20 HIF-1-dependent muscle DEGs with evolutionarily conserved homologs and available genetic tools. Specifically, we performed a muscle-specific RNAi screen for modulators of sensitivity to levamisole (Extended Data Fig. 23c,d). *tspo-1*, which encodes the evolutionarily conserved translocator protein TSPO, emerged as a candidate. Despite being broadly expressed across muscle subtypes, *tspo-1* displayed HIF-1-dependent downregulation specific to vulval muscle cluster 4 (Fig. 4f; Extended Data Fig. 23e). Like *egl-9* mutants, *tspo-1* null mutants exhibited hypersensitivity to levamisole (Extended Data Fig. 23f). Expression of wild-type *tspo-1* in muscle using the *myo-3* promoter rescued the levamisole hypersensitivity of *tspo-1* mutant animals, indicating that *tspo-1* functions in muscle to regulate levamisole sensitivity (Extended Data Fig. 23f). Consistent with a subtype-specific contribution, expression of *tspo-1(+)* in cluster-4/vulval muscle cells using either the *srb-16* (Extended Data Fig. 24) or *unc-103e* promoter rescued the levamisole hypersensitivity, whereas expression in cluster-8 body-wall muscle cells (in which *tspo-1* is not HIF-1-regulated) did not rescue (Fig. 4g). Thus, *tspo-1* function in vulval muscle cells, but not in cluster 8 body-wall muscle cells, is sufficient to restore normal levamisole responses in *tspo-1* mutants. To determine whether HIF-1-dependent regulation of TSPO might be evolutionarily conserved, we examined TSPO levels in human AC16 cardiomyocytes following HIF-1α activation. Treatment with cobalt chloride, a chemical inducer of HIF-1α^40^, reduced TSPO levels, and this reduction was suppressed by HIF-1α knockdown (Fig. 4h,i; Extended Data Fig. 25), indicating that HIF-1-dependent regulation of TSPO might be evolutionarily conserved. Together, these findings provide functional validation of our identification of a cell-subtype-specific HIF-1-responsive gene.

## Discussion

By analyzing patterns of gene expression at the single-cell level in an intact animal, we observed that HIF-1 activation modulates a shared central-carbon-metabolism core together with largely both tissue- and cell-type-dependent transcriptional responses. Two complementary analytical approaches support this conclusion. First, a standard analysis of differential gene expression showed that the identities of HIF-1-dependent genes differed extensively among neurons, intestine and muscle and among selected cell subtypes within these tissues, indicating that the tissue differences reflect distinct sets of responsive genes rather than only quantitative scaling of a single program. Supervised machine learning provided a complementary multivariate view that did not depend on the requisite significance thresholds of differential-expression analysis: genotype-defined HIF-1 states could be distinguished from single-cell transcriptional profiles within each tissue, and the genes most informative for prediction showed limited overlap among tissues, consistent with the tissue-contextualized responses identified by differential-expression analysis. Strikingly, the predictive-gene rankings overlapped only partly with cluster-wise differential-expression results, indicating that the multivariate analysis identified predictive information that did not meet cluster-wise differential-expression thresholds. Together, these analyses indicate that the shared central-carbon-metabolism response is embedded within broader, largely tissue-specific transcriptional programs. The context dependence of the effects of HIF-1 activation extended below the level of whole tissues to anatomically and functionally specialized cell subtypes. Selected neuronal, intestinal and muscle subtypes shared some enriched functional categories but also showed subtype-associated enrichment patterns. HCR smFISH and transcriptional-reporter analyses validated the identification of the predominant anatomical sites of HIF-1-dependent induction predicted by scRNA-Seq. The identification of *tspo-1* provides a functional example of a cell subtype-specific and possibly evolutionarily conserved HIF-1-responsive gene identified through the atlas and illustrates how the atlas can identify cell-subtype-specific HIF-1-responsive genes that would not be apparent from measurements of gene expression in whole animals.

The mechanisms that generate these context-dependent responses remain unclear. Nearly all genes induced across all three tissues showed evidence consistent with direct HIF-1 binding, whereas tissue-specific upregulated genes included genes both with and without detectable HIF-1 occupancy. These patterns are consistent with a greater contribution from direct HIF-1 regulation to the shared transcriptional core, whereas tissue-specific responses likely involve a combination of context-dependent direct regulation and downstream indirect mechanisms. HIF-1 binding was substantially less frequent among downregulated genes, and most if not all HIF-1 downregulation is likely indirect. A complementary perspective comes from our supervised-learning analyses: predictive-gene rankings overlapped in part with differential-expression results for individual clusters; however, many genes that did not meet cluster-wise differential-expression thresholds also were major contributors to the prediction of genotype-defined HIF-1 states. We speculate that relatively small changes in the levels of expression of some HIF-1-responsive genes collectively contribute to the transcriptional representation and impact of HIF-1 activity. Differences in chromatin accessibility, cis-regulatory architecture, co-regulator availability and signaling inputs might contribute to determining which genes respond to HIF-1 in each cellular context. More generally, our analyses suggest that examining a holistic cellular transcriptome might better inform our understanding of the activity of a transcription factor than would using differential transcriptomics alone. We suggest that machine-learning multivariate analyses similar to those we performed could increase our fundamental understanding of the activities of transcription factors other than HIF-1 as well as of the consequences on gene expression of such processes as intercellular signaling. We hope that our atlas and analytical framework provide both a resource and a roadmap for identifying regulatory mechanisms and physiological effectors that shape tissue- and cell-type-specific responses to hypoxia and to other environmental and physiological stimuli.

### Limitations

Several limitations should be considered when interpreting the machine-learning analyses. Each tissue-genotype dataset was generated from a separate preparation containing cells pooled from many animals. Although pooling reduces sensitivity to variation among individual animals, genotype could not be fully separated from preparation of origin because independent replicate preparations were not profiled for each tissue-genotype combination. Accordingly, cell-level cross-validation evaluates prediction across held-out cells from the available pooled preparations rather than generalization to independently prepared biological samples. The high discriminatory performance could therefore reflect a combination of HIF-1-dependent transcriptional biology and preparation-specific variation. Several observations support a substantial biological component. Genes repeatedly recovered among the informative features included established HIF-1-responsive genes, such as *pck-1*, *cysl-2* and *F22B5.4*, whose expression increased in *egl-9* mutants and reversed in *egl-9 hif-1* double mutants, matching the expected direction of HIF-1-dependent regulation. Predictive genes also overlapped across the two genotype contrasts in neurons and muscle. Nevertheless, these observations do not eliminate sample-of-origin or other preparation-specific confounding.

Permutation-importance results should also be interpreted as measures of predictive contribution rather than definitive rankings of biological importance. Feature importance was estimated from a single 80/20 evaluation split after model selection, and the selected model– transformation configuration differed among tissues and genotype comparisons. The standard deviation across repeated permutations reflects variability of the permutation procedure within the same evaluation subset and does not measure sensitivity to the choice of train/evaluation split. Exact gene rankings can therefore depend on the specific data split, model, preprocessing representation and correlations among gene-expression features. Correlated genes might carry redundant predictive information, such that permuting one feature can underestimate its contribution when related features remain available to the model. This limitation is particularly relevant when baseline classification performance approaches the AUROC ceiling, where permutation of individual genes can produce relatively small changes in performance. In the intestinal analyses, extensive ties and variability in permutation-importance values around the rank cutoffs further make the exact membership of the top-gene sets sensitive to small differences in rank. We therefore interpret individual permutation-importance values, exact gene ranks and cross-tissue or cross-contrast overlaps as exploratory measures of predictive contribution rather than evidence of causal, independent or biomarker-level importance.

## Acknowledgements

We thank members of the MIT BioMicro Center for technical support with scRNA-Seq experiments; WormBase; the *Caenorhabditis* Genetics Center and the National BioResource Project for *C. elegans* strains; the ENCODE Consortium, members of the Reinke laboratory for generating *C. elegans* ChIP-Seq data of the modERN project; Na An for critical operational support; other members of the Horvitz laboratory, especially Calista Diehl, for technical support, advice, and suggestions about the manuscript; and Dongyeop Lee and Wenhan Chang for helping generate reagents. Cartoon figures were created with BioRender.com. D.D.G. was supported by the Sara Elizabeth OʹBrien Trust, Bank of America, N.A., Co-Trustees. This work was supported in part by the MIT Koch Institute Support (core) Grant P30-CA14051 from the National Cancer Institute and by the MIT McGovern Institute for Brain Research. This work was also supported by the National Institutes of Health (R01GM024663) and the Howard Hughes Medical Institute. H.R.H. is an investigator of the Howard Hughes Medical Institute.

## Author contributions

J.N.K. and H.R.H. conceptualized the study. J.N.K., D.D.G., and H.R.H. designed experiments. J.N.K. conducted the scRNA-Seq experimental studies. J.N.K., D.D.G., D.M., and S.R.S. analyzed scRNA-Seq datasets. J.N.K. and L.G. conducted HCR-FISH experiments. I.K. generated transcriptional reporters. J.N.K. and D.M. performed and analyzed machine-learning benchmarking analyses. D.M. developed the machine-learning script. J.N.K., D.D.G., and A.S. conducted the behavioral assays. J.N.K. conducted expression pattern analyses. J.N.K. conducted human cell culture experiments. J.N.K. and H.R.H. wrote the paper with input from D.D.G. and other authors.

## Data and materials availability

The single-cell RNA sequencing (scRNA-Seq) dataset generated and analyzed during this study is available in the Gene Expression Omnibus (GEO) repository under accession number GSE264617.

## Materials and Methods

### Strains

*C. elegans* hermaphrodite strains were maintained at 20 °C as previously described^41^. The N2 Bristol strain was the reference wild-type strain. All strains were derived from Bristol N2. Standard molecular biology and microinjection methods, as previously described^42^, were used to generate transgenic worms. The transgenes and specific mutations used are listed below:

LG I: *ccIs4251[P_myo-3_::GFP(NLS)::LacZ (pSAK2) + P_myo-3_::GFP (mitochondrially targeted) (pSAK4) + dpy-20(+)];* LG III: *otIs173[P_rgef-1_::DsRed2 + ttx-3promB::GFP], unc-119(ed3), oxTi872[P_vha-6_::GFP::tbb-2 3’UTR + Cbr-unc-119(+)];* LG V: *egl-9(sa307), hif-1(ia4)*.

Extrachromosomal array:

*nEx3267-8[P_srb-16_::GFP + P_unc-103e_::BFP + P_myo-3_::mCherry], nEx3226-7[P_C36C5.5_::GFP + P_nmy-3_::BFP + P_myo-3_::mCherry],*

Strain identifiers, including genotype, of all mutants and transgenic animals used in this study are provided in Extended Data Table 6.

### Single-cell RNA-sequencing

#### Dissociation of animals into cell suspensions

Single-cell suspensions were generated as described^18^ with minor modifications. Briefly, synchronized populations of worms were grown on NGM plates seeded with OP50 to the stage of young (one-day) adults. Worms were harvested from these plates, washed three times with ice-cold M9 buffer and treated with SDS-DTT (200 mM DTT, 0.25% SDS, 20 mM HEPES, 3% sucrose, pH 8.0) for 6 min. Worms then were washed five times with egg buffer (118 mM NaCl, 48 mM KCl, 2 mM CaCl_2_, 2 mM MgCl_2_, 25 mM HEPES, 340 mOsm, pH 7.3) and treated with pronase (15 mg/mL) for 15-20 min. During pronase treatment, worm suspensions were pipetted up and down rapidly in four sets of 100 repetitions each. Pronase treatment was stopped by the addition of ice-cold L-15–10 media (90% L-15 media, 10% FBS). The suspension was then passed through a 35 μm nylon filter into a collection tube, washed once with 1x PBS, and prepared for FACS.

#### FACS of fluorescently-labeled neuronal, intestinal and muscle cells

FACS was performed using a BD FACS Aria III cell sorter running BD FACS Diva software (BD FACSARIA III cell sorter, RRID:SCR_016695). DAPI was added to samples at a final concentration of 1 μg/mL to identify membrane-compromised cells and exclude non-viable cells during sorting. DsRed-positive DAPI-negative neurons, GFP-positive DAPI-negative intestinal cells, or GFP-positive DAPI-negative muscle cells were sorted from the single-cell suspension into 1x PBS containing 1% FBS. Non-fluorescent and single-color controls were used to set gating parameters. Cells were then concentrated and processed for single-cell sequencing.

#### Single-cell RNA sequencing and data processing

Samples were processed for single-cell sequencing using the 10X Genomics Chromium 3’mRNA-sequencing platform. Libraries were prepared using the Chromium Next GEM Single Cell 3’ Kit v3.1 according to the manufacturer’s protocol. The libraries were sequenced using an Illumina NextSeq 500 with 75 bp paired end reads.

Mapping and counting were carried out using 10X Genomics’ CellRanger software/5.0.0. The sparse matrices in .mtx format were further processed using Seurat 4.0.2 based on r/4.0.4. Neuron samples, muscle cell samples, and intestinal cell samples were processed separately. Genes expressed in fewer than 5 cells were filtered out, and cells with at least 200 detected genes were retained for downstream analyses. Expression counts were normalized using Seurat’s “LogNormalize” method, and the top 2,000 highly variable genes were identified using the FindVariableFeatures(method = “vst”) function for downstream dimensionality reduction and clustering analyses. Data from the three different genotypes were integrated using shared correlation anchors. Then a linear transformation step was applied by scaling the expression per gene to be mean 0 and variance to be 1. Principal component analysis (PCA) was performed on the variable genes using the RunPCA function in Seurat. UMAPs were created using the top 30 principal components. Cluster-specific marker genes were identified using Seurat’s FindAllMarkers function, and cluster identities were assigned based on expression of established cell-type-specific marker genes. Differentially expressed genes across genotypes within each cluster were identified using Seurat’s FindMarkers function. Expression patterns of selected differentially expressed genes were visualized across clusters using feature plots and violin plots to assess consistency of differential expression patterns across samples.

Power calculations were performed using the Seurat power calculator.

#### Hypoxia Response Element (HRE) Motif Search

The 1,000 bp upstream sequences of differentially regulated genes were identified using the getfasta command from BEDTools (v2.29.2). HRE motifs—specifically GCGTG and ACGTG^22^ —were identified using custom scripts developed by the MIT BioMicro Center. For genes located on the reverse strand, the reverse complements of the HRE motifs (CACGC and CACGT) were sought instead. The genomic coordinates of the identified HRE motifs and their distances from the transcription start site were annotated using additional in-house tools.

#### HIF-1 Binding Site Identification

HIF-1 binding peaks were obtained from the modERN HIF-1 ChIP-Seq dataset, in which HIF-1 occupancy was mapped using a GFP-tagged HIF-1 protein and an anti-GFP antibody (strains OR3349 and OR3350; ENCODE accessions ENCSR991HIA and ENCSR517OZM; Vora *et al.* (2022)), using the peak file (modENCODEpeaks.txt) downloaded from the ENCODE portal. A differentially expressed gene was scored as HIF-1-bound if a ChIP-Seq peak overlapped or was proximal to the gene, using custom tools developed by the MIT BioMicro Center.

#### Hybridization chain reaction (HCR) single-molecule fluorescence *in situ* hybridization (smFISH) Probe Design and Staining Protocol

Custom HCR v3.0 probe sets targeting *gcy-33*, T10B9*.9*, *ftn-1*, *mtl-1*, *mmcm-1*, *oac-*54, *pcca-1,* and *pccb-1* designed based on mRNA sequences from the NCBI Reference Sequence (RefSeq) database, were obtained from Molecular Instruments. Each probe set was synthesized using 20 split-initiator probe pairs.

HCR staining^43^ was performed using the Molecular Instruments HCR RNA-FISH protocol for *C. elegans* larvae (Rev. 9, 2023), with modifications to optimize staining for young adult animals.

Briefly, fixed whole-mount worms were permeabilized using 1% β-mercaptoethanol and 100ug/mL proteinase K, followed by overnight probe hybridization at 37 °C. After post-hybridization washes, amplification was performed using snap-cooled hairpins (H1 and H2) incubated overnight at room temperature in the dark. Samples were subsequently washed, counterstained with DAPI, and stored at 4 °C in the dark prior to imaging. Detailed protocol steps are provided in the Supplementary Methods.

#### Gene ontology (GO) enrichment analysis

To generate diagrams containing counts of genes up- or downregulated in each tissue, the numbers of genes that were either up- or downregulated significantly (adjusted p<0.05) by more than 2-fold in *egl-9* compared to wild-type in at least one cluster of the tissue were counted.

Those genes that were also down- or upregulated by more than 2-fold in *egl-9 hif-1* compared to *egl-9* in the same cluster were considered *hif-1*-dependent. This analysis was performed using Python with the Pandas v1.5.3 package for dataframe manipulation.

The list of genes found to be differentially up- or downregulated in an EGL-9/HIF-1 dependent manner by more than 2-fold in any cluster were used for gene ontology (GO) enrichment analysis. The PANTHER Overrepresentation Test (Released 20231017) was used to query the gene list against the GO Ontology database (DOI: 10.5281/zenodo.7942786. Released 2023-01-05). Enriched GO terms were determined by Fisher’s exact test with false discovery rate correction. The figure was created using drawsvg v2.3.0 and Python v3.12.0 scripts.

#### Machine learning framework for classifying HIF-1 activation states from single-cell transcriptomes

To identify gene expression patterns associated with HIF-1 activation, we applied a GPU-accelerated supervised machine-learning framework to single-cell transcriptomic profiles using PyTorch and XGBoost. The expression levels of genes in individual cells served as input features, and genotype served as the binary class label for each comparison. Neuronal, intestinal, and muscle cells were analyzed separately. For each tissue, we performed two complementary binary genotype comparisons—wild type versus *egl-9* and *egl-9* versus *egl-9 hif-1*— to assess transcriptional states associated with HIF-1 activation and their HIF-1 dependence, yielding six classification tasks in total. The goal was to determine whether transcriptomic information could distinguish genotype-defined HIF-1 activation states and to identify genes that contributed to predictive performance.

#### Data preprocessing and gene selection

The machine-learning (ML) analysis used RNA-assay expression values generated using Seurat LogNormalize, with normalization performed separately for each sample before integration; no integrated or batch-corrected expression values were used as model input. The resulting expression matrices were organized with genes as rows and individual cells as columns. The matrices were transposed for model training so that rows represented individual cells and columns represented genes. Each cell was assigned a binary class label according to genotype, using genotype information encoded in the cell identifier.

Within each cross-validation training fold, genes were retained if (i) their maximum normalized expression across training cells exceeded 0.1 and (ii) the ratio of maximum to minimum expression across training cells exceeded threefold. Minimum expression values below 0.001 were set to 0.001 before calculation of this ratio to avoid division by zero or near-zero values. This filtering step was used to reduce dimensionality by excluding genes with negligible expression or limited variation across the training cells. Gene selection was performed using the training cells only within each cross-validation fold, and the resulting feature set was subsequently applied to the corresponding held-out cells.

To evaluate whether predictive performance depended on data representation, we compared three forms of the normalized expression matrix: (i) the original normalized expression values without an additional transformation; (ii) square-root-transformed values, which reduced the influence of large expression values; and (iii) rank-based quantile-transformed values, which emphasized the relative ordering of expression values across cells for each gene rather than their absolute magnitudes. Rank-based quantile transformation mapped expression values to a uniform distribution and was fitted using the training cells only within each cross-validation fold before being applied to the corresponding held-out cells.

Within each cross-validation fold, expression values for each gene were further standardized by subtracting the mean and dividing by the standard deviation calculated from the training cells. The same training-derived scaling parameters were then applied to the corresponding held-out cells. Thus, gene selection, rank-transformation fitting, and feature standardization during cross-validation were determined using the training partition only.

#### Model training and evaluation

We compared nine supervised machine-learning approaches for distinguishing genotype-defined HIF-1 activation states from gene-expression profiles. These included an unregularized logistic classifier and three regularized linear classifiers: ridge classification with an L2 penalty, lasso classification with an L1 penalty, and elastic-net classification combining L1 and L2 penalties. These linear approaches classify cells using weighted combinations of gene-expression features; regularization limits model complexity and reduces the influence of uninformative or redundant features. We also evaluated k-nearest-neighbor classification, which classifies each cell according to its most transcriptionally similar cells; a multilayer neural network, which can model nonlinear relationships among genes; and three tree-based ensemble approaches— implemented as random-forest-style, extra-trees-style, and gradient-boosting models—which combine predictions from many decision trees. Evaluating multiple model classes allowed predictive performance to be compared across modeling approaches rather than relying on a single algorithm.

The logistic, regularized linear, and neural-network classifiers were implemented in PyTorch and trained using binary cross-entropy loss and the Adam optimizer. The neural network contained two hidden layers with 100 and 50 units, rectified linear-unit activation functions, and L2 regularization. The k-nearest-neighbor classifier used five nearest neighbors and brute-force GPU distance calculations. The tree-based ensemble models were implemented using XGBoost with GPU histogram acceleration. The random-forest-style model contained 200 trees with a maximum depth of eight, no learning-rate shrinkage, and 80% row and feature subsampling. The extra-trees-style model used additional feature subsampling at individual tree nodes. The gradient-boosting model contained 100 trees with a maximum depth of three and a learning rate of 0.1.

Each model–transformation combination was evaluated using five-fold stratified cross-validation. Cells were divided into five folds while maintaining similar proportions of the two genotype classes in each fold. During each iteration, four folds, representing approximately 80% of the cells, were used for model training, whereas the remaining fold was held out for evaluation. This process was repeated five times, with each fold serving as the held-out fold once. Thus, every cell was evaluated by a model that had not been trained on that cell. Gene filtering, rank-transformation fitting, and standardization were performed independently within each cross-validation iteration using the training cells only.

Model performance was assessed using the area under the receiver-operating-characteristic curve (AUROC), which quantifies discrimination between the two genotype classes across classification thresholds. An AUROC of 0.5 indicates chance-level discrimination, whereas an AUROC of 1 indicates perfect discrimination. AUROC was calculated separately for each of the five held-out folds. The five fold-level AUROC values were then averaged to obtain the mean cross-validated AUROC for each model–transformation combination, and the standard deviation across the five folds was calculated as a measure of fold-to-fold variability. For each tissue and genotype comparison, a model–transformation configuration with the highest mean cross-validated AUROC was selected for subsequent permutation-importance analysis; where configurations were tied, one of the tied configurations was selected.

#### Identification and comparison of predictive genes

To identify genes contributing to classification performance, we performed permutation-importance analysis using the model–transformation configuration selected for each tissue and genotype comparison. The selected configurations differed among tissues and genotype comparisons and are listed in Extended Data Table 3. After model selection by five-fold cross-validation, a separate stratified 80/20 train/evaluation split of the full dataset was generated using a fixed random seed of 42. The selected configuration was fitted using the 80% training subset, and permutation importance was evaluated using the corresponding 20% evaluation subset. Gene filtering, transformation fitting where applicable, and z-score standardization were determined using only the 80% training subset and then applied to the 20% evaluation subset. In these permutation-importance analyses, the retained feature sets comprised 16,304 neuronal, 11,557 muscle, and 10,569 intestinal genes for the wild-type versus *egl-9* comparison, and 16,239 neuronal, 11,863 muscle, and 10,119 intestinal genes for the *egl-9* versus *egl-9 hif-1* comparison.

For each gene, its expression values were randomly permuted across cells in the 20% evaluation subset while the values of all other genes were left unchanged, and AUROC was recalculated. This procedure was repeated 30 times for each gene. Gene importance was quantified as the mean decrease in AUROC following permutation relative to the baseline AUROC of the unpermuted evaluation data. The standard deviation across the 30 permutations was also calculated; this value reflects variability of the permutation procedure within the same evaluation subset and does not measure sensitivity to the choice of train/evaluation split.

Genes whose permutation produced larger decreases in AUROC were interpreted as making stronger contributions to predictive performance and were ranked according to their mean permutation-importance values. We refer to these genes as informative or predictive genes rather than biomarkers because permutation importance measures contribution to model prediction and does not establish biological causality, independent necessity, or diagnostic utility. In addition, because correlated or co-expressed genes can carry redundant predictive information, permutation of one feature may underestimate its contribution when related features remain available to the model. Individual importance values and exact gene ranks were therefore interpreted as measures of predictive contribution rather than definitive rankings of biological importance.

For comparisons between the two genotype contrasts within a tissue, expected overlap under random selection was calculated accounting for the numbers of genes eligible for ranking in each contrast and the number shared between the two retained-gene backgrounds. For two independently selected sets of size *k*, drawn from backgrounds containing *N*₁ and *N*₂ genes with *M* genes shared between them, the expected intersection size was *E[I]* = *Mk*²/(*N*₁*N*₂). The corresponding reference Jaccard value was calculated as *E[I]*/[2*k* − *E[I]*]. The retained-gene backgrounds contained 16,304 and 16,239 genes in neurons, 10,569 and 10,119 genes in intestine, and 11,557 and 11,863 genes in muscle; the corresponding numbers of genes shared between the two backgrounds were 16,078, 10,045 and 11,412, respectively.

For heatmap visualization, permutation-importance values were min–max normalized separately within the top 50 genes for each tissue and genotype comparison: normalized importance = [importance − minimum(top 50)]/[maximum(top 50) − minimum(top 50)]. Thus, the highest- and lowest-importance genes within each top-50 list were assigned normalized values of 1 and 0, respectively. Because normalization was performed separately for each tissue and genotype comparison, these values represent relative importance within each analysis and were not used to compare absolute permutation-importance magnitudes across tissues or genotype contrasts.

#### Behavioral assay: Levamisole sensitivity

The day before the assay, 100 μl of an overnight inoculated OP50 was spread onto 60 mm NGM plates and incubated at 37°C overnight. On the following day, plates were dried under a hood for 30 min to remove excess moisture. Levamisole (Millipore Sigma L9756) solutions of the specified concentration were prepared using M9 buffer. Approximately 10 adult *C. elegans* worms were picked and gently washed from the pick onto the assay plate with 10 μl of the levamisole solution. Mobility was scored after 10 min of levamisole exposure by assessing whether animals moved at least one full body length within a 30 sec observation period. Assay plates were gently tapped to confirm that animals retained the ability to move.

#### RNAi treatment

RNAi by feeding was performed as described previously^44,45^. In brief, HT115 *E. coli* bacteria carrying RNAi clones in the pL4440 vector were grown for at least 12 hr in Luria broth (LB) liquid medium with 75 mg/L ampicillin at 37°C. These cultures were seeded onto 60 mm plates with NGM containing 1 mM isopropyl-β-D-thiogalactopyranoside (IPTG) (Amresco) and 75 mg/L ampicillin and incubated for 24 hr at 37°C. For the levamisole assays, three adult animals were added to each RNAi plate and their progeny were used to test levamisole sensitivity as described above. Each RNAi experiment was performed alongside an empty pL4440 vector negative control.

#### Microscopy

Confocal images were obtained using an LSM 800 confocal microscope (Zeiss LSM 800 with Airyscan Microscope, RRID:SCR_015963) and ZEN software. Confocal images of *C. elegans* were obtained using 10x air and 63x oil objectives (Zeiss). Images were processed and prepared for publication using FIJI software (Fiji, RRID:SCR_002285) and Adobe Illustrator (Adobe Illustrator, RRID:SCR_010279).

#### Cultivation and treatment of AC16 cells

The AC16 human cardiomyocyte cell line (Millipore Sigma SCC109) was cultured in Dulbecco’s Modified Eagle’s Medium: Nutrient Mixture F-12 (DMEM / F-12) (ThermoFisher Scientific 11330032), supplemented with 10% fetal bovine serum (Gibco 10437) and 1% penicillin-streptomycin (Gibco 15140122). Cells were maintained in a humidified incubator at 37°C with 5% CO_2_. For experiments, cells were seeded about 50,000 cells/cm^2^ density in DMEM growth medium and treated for 24 hr with cobalt (II) chloride (Millipore Sigma C8661) at the specified concentrations.

#### siRNA transfection

ON-TARGETplus Human HIF-1α siRNA (Dharmacon L-004018-00-0005), ON-TARGETplus Non-targeting Control Pool (Dharmacon D-001810-10-05), and siGLO RISC-free control siRNA (Dharmacon D-001600-01-05) were purchased from Dharmacon. Using Lipofectamine RNAiMAX (ThermoFisher 13778075), cells were transfected with either HIF-1α siRNA SMARTpool reagent (50 nM), or non-targeting control siRNA (50 nM). siGLO RISC-free control siRNA (5 nM) was co-transfected in all samples. Cells were fixed after 48 hr and further processed for immunocytochemistry.

#### Immunocytochemistry

AC16 cells were grown on Nunc Lab-Tek II Chamber slides (ThermoFisher, 154461). Cells were fixed with 4% paraformaldehyde (PFA) / PBS for 15 min and then permeabilized by incubation with 0.2% Triton X-100 in PBS for 10 min at room temperature. Nonspecific binding sites were blocked with 3% BSA/PBS for 1 hr at room temperature. Cells were then subsequently stained with primary antibodies against (HIF-1α antibody, Cell Signaling 36169; TSPO antibody, ThermoFisher MA5-24844). Secondary antibodies Alexa Fluor 594 donkey anti-rabbit IgG (H+L) (ThermoFisher A21207) or Alexa Fluor 488 goat anti-rabbit IgG (H+L) (ThermoFisher A11034) were diluted 1:2000 in 0.1% ovalbumin/PBS and incubated with the samples for 1 hr at room temperature. After washing, slides were mounted using SlowFade diamond antifade mountant with DAPI (ThermoFisher S36964) to visualize nuclei. Confocal images of cells were obtained using 10x air and 63x oil objectives lens (Zeiss) on a Zeiss LSM800 confocal microscope. Images were processed with ImageJ software (NIH) software.

#### Quantification of mammalian cell staining

Randomly chosen sections of images (blinded) were analyzed using Fiji/Image J software. In brief, the Identify Primary Objects, Measure Object Intensity, and Export to Spreadsheet modules were used sequentially for nuclear segmentation (DAPI channel), nuclear intensity measurements (HIF-1α), cytosol intensity measurements (TSPO), and data export, respectively.

#### Statistical analysis

For all other experiments, one-way ANOVA was used to assess statistical differences. Post-hoc Tukey-Kramer tests were conducted for pairwise comparisons between all conditions, while post-hoc Bonferroni or Dunnett tests as appropriate were performed for comparisons to control. Significance levels were denoted as follows: * for p < 0.05, ** for p < 0.01, and *** for p < 0.001. Statistical analyses were conducted using GraphPad Prism software (version 10.0.3 (217)).

---

**Extended Data Figure 1.**
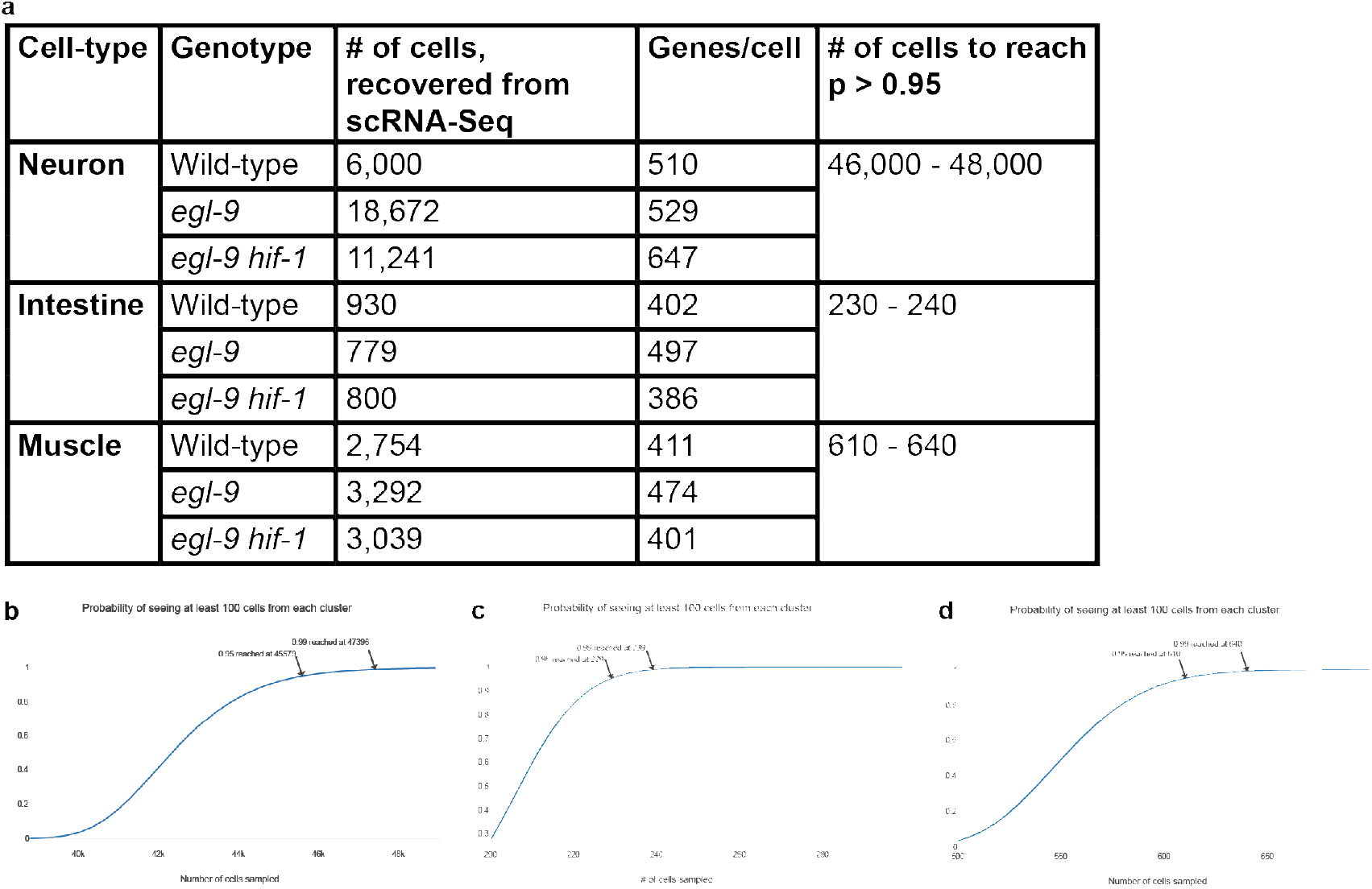
Power analysis for minimum cells required for scRNA-Seq analyses. **a**, Number of cells recovered from scRNA-Seq experiment pipeline, average number of genes detected per cell, and number of cells required for p > 0.95. **b-d,** Power analyses of cells required for comprehensive identification of the (**b**) ∼120 expected cell types within neuronal tissue, (**c**) ∼2 expected cell types within intestinal tissue, and (**d**) ∼20 expected cell types within muscle tissue^13,14,46,47^. Complete neuronal identification from *C. elegans* scRNA-Seq analyses has been accomplished previously by others, first for embryonic and then for adult neurons^18,48^. Based on our power calculation^25,27,49,50^, about 47,000 neuronal cells would be required to identify all neuronal subtypes (b). Our dataset falls below this threshold and therefore does not capture all neuronal classes, though it is sufficient to resolve major subpopulations. (c,d), The numbers of intestinal and muscle cells analyzed are expected to be sufficient to identify all major intestinal and muscle subtypes.

**Extended Data Figure 2.**
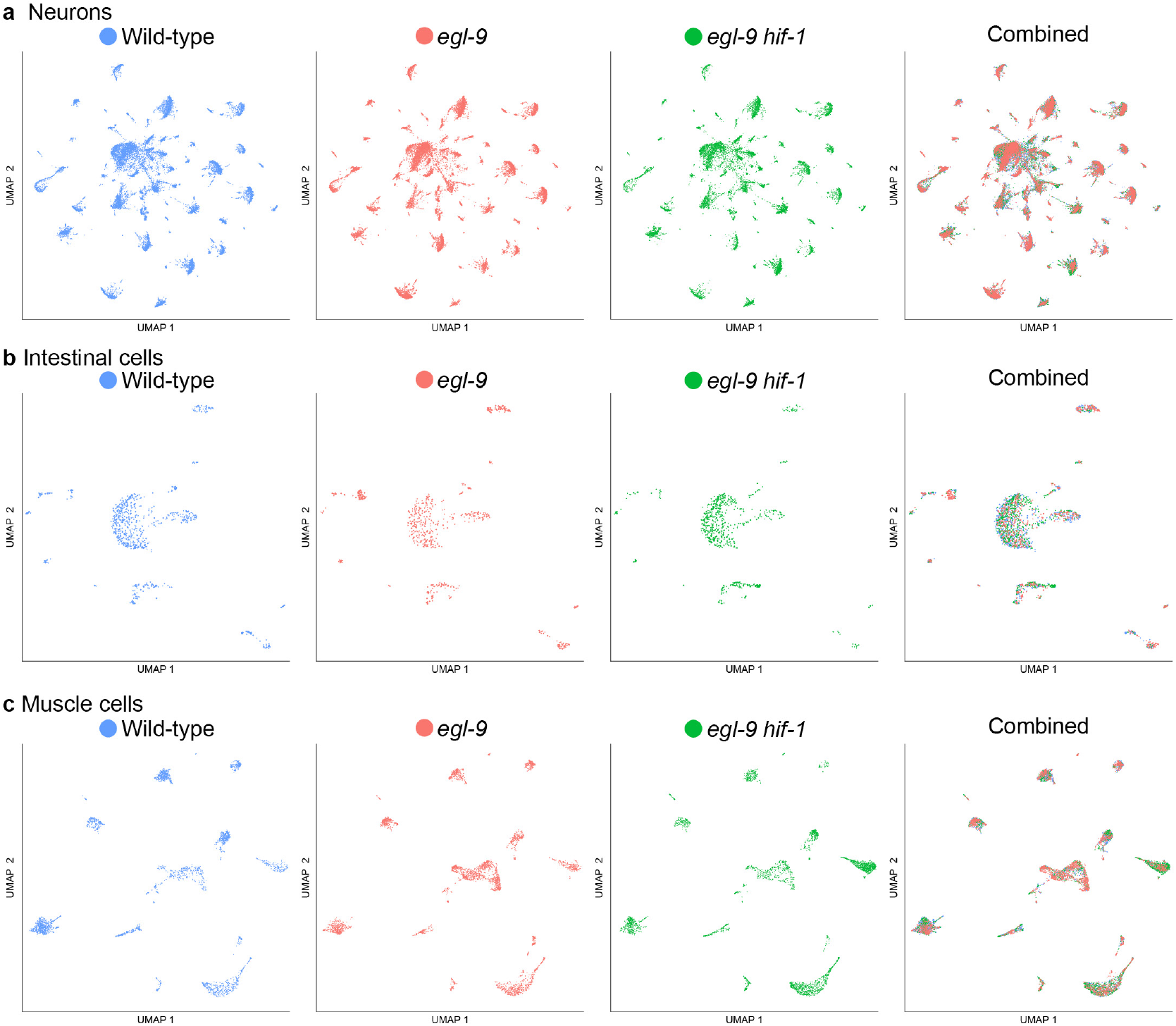
UMAPs of neuronal, intestinal, and muscle cells by genotype. **a**, UMAP (Uniform Manifold Approximation and Projection) projections of neurons separated by genotype. Blue, red, and green represent wild-type, *egl-9* mutant, and *egl-9 hif-1* double-mutant cells, respectively. **b,** UMAPs of intestinal cells separated by genotype. **c,** UMAPs of muscle cells separated by genotype.

**Extended Data Figure 3.**
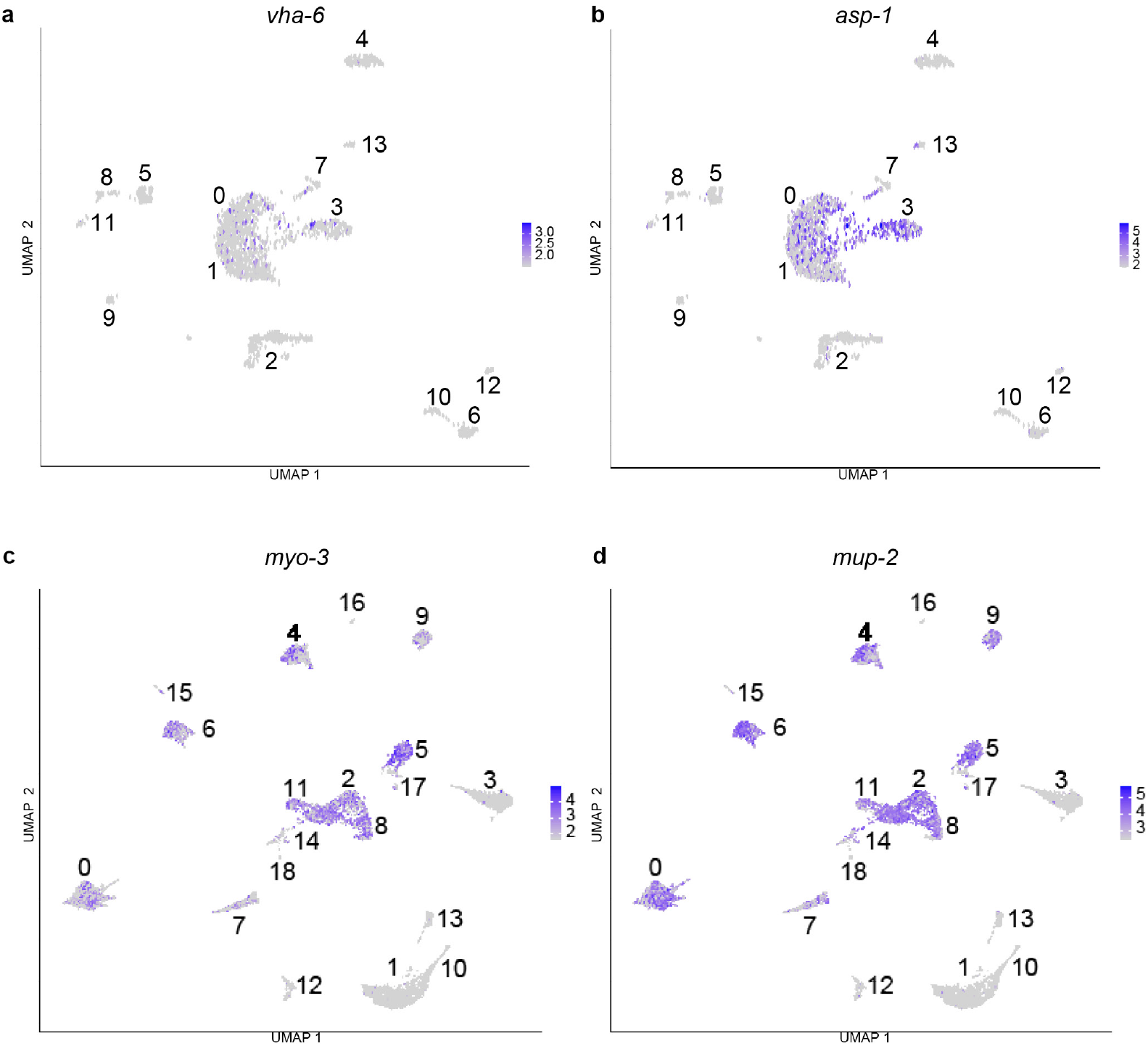
Identification of intestinal and muscle cell clusters. **a,b**, UMAP projection of cells sorted using the intestinal marker *vha-6* overlaid with expression of intestinal markers (**a**) *vha-6* and (**b**) *asp-1*. UMAP clusters 0 and 3 represent intestinal cell clusters that express these previously established markers. **c,d,** UMAP projection of cells sorted using the muscle marker *myo-3* overlaid with expression of muscle markers (**c**) *myo-3* and (**d**) *mup-2*. UMAP clusters 0, 2, 4, 5, 6, 7, 8, 9, and 11 represent muscle cell clusters that express these previously established markers.

**Extended Data Figure 4.**
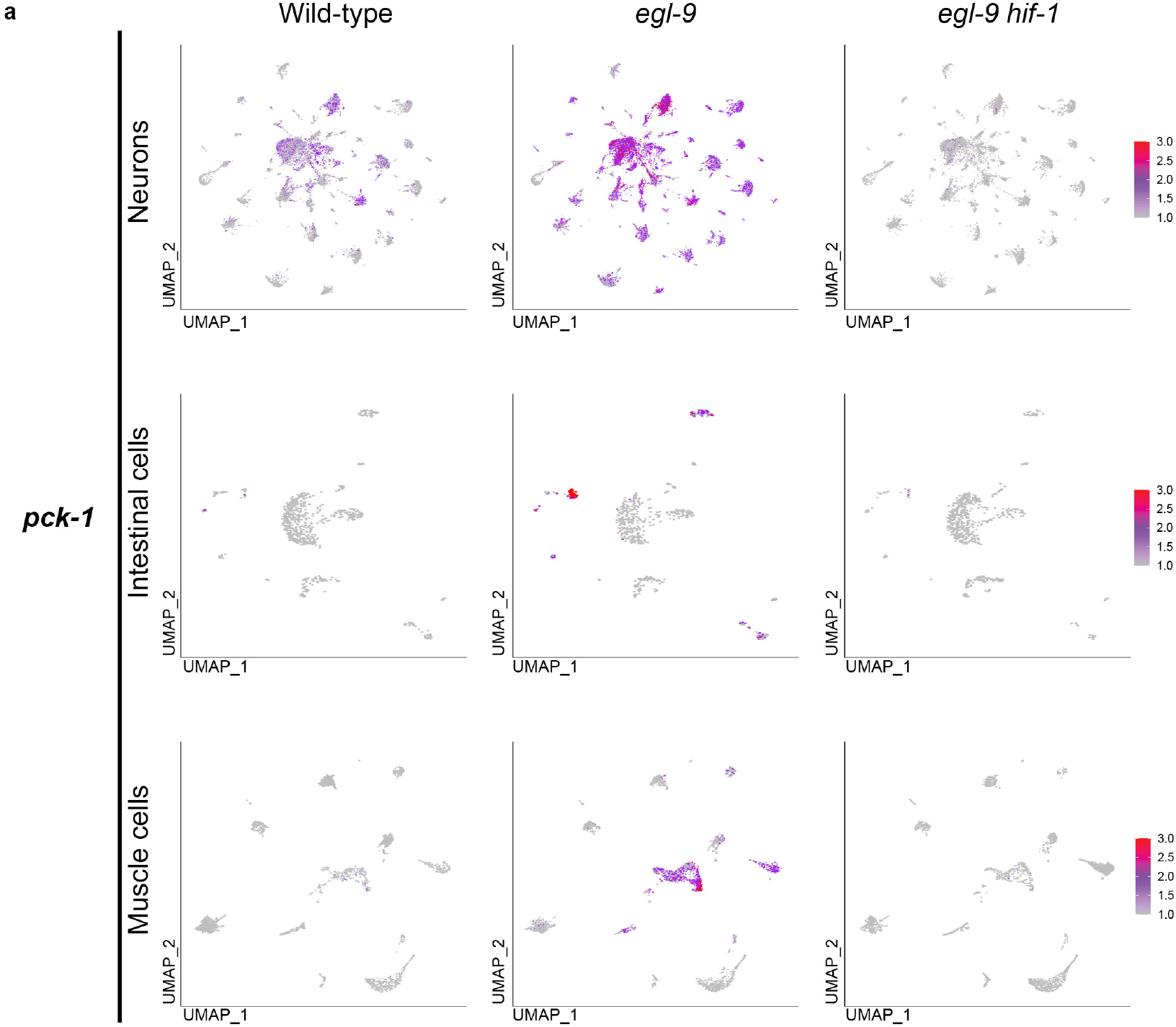
Visualization of *pck-1* expression in UMAPs of neuronal, intestinal, and muscle tissue by genotype. **a**, UMAPs of neuronal, intestinal, and muscle tissue separated by genotype showing differential expression of *pck-1* – a gene known to be activated by HIF-1^22^ – in wild-type, *egl-9* mutant, and *egl-9 hif-1* double-mutant animals. Each projection shows the color-coded expression levels of *pck-1* in cells from that genotype. As expected, *egl-9* mutants exhibited significantly increased expression of *pck-1* in numerous cell types compared to wild-type or *egl-9 hif-1* double-mutant animals.

**Extended Data Figure 5.**
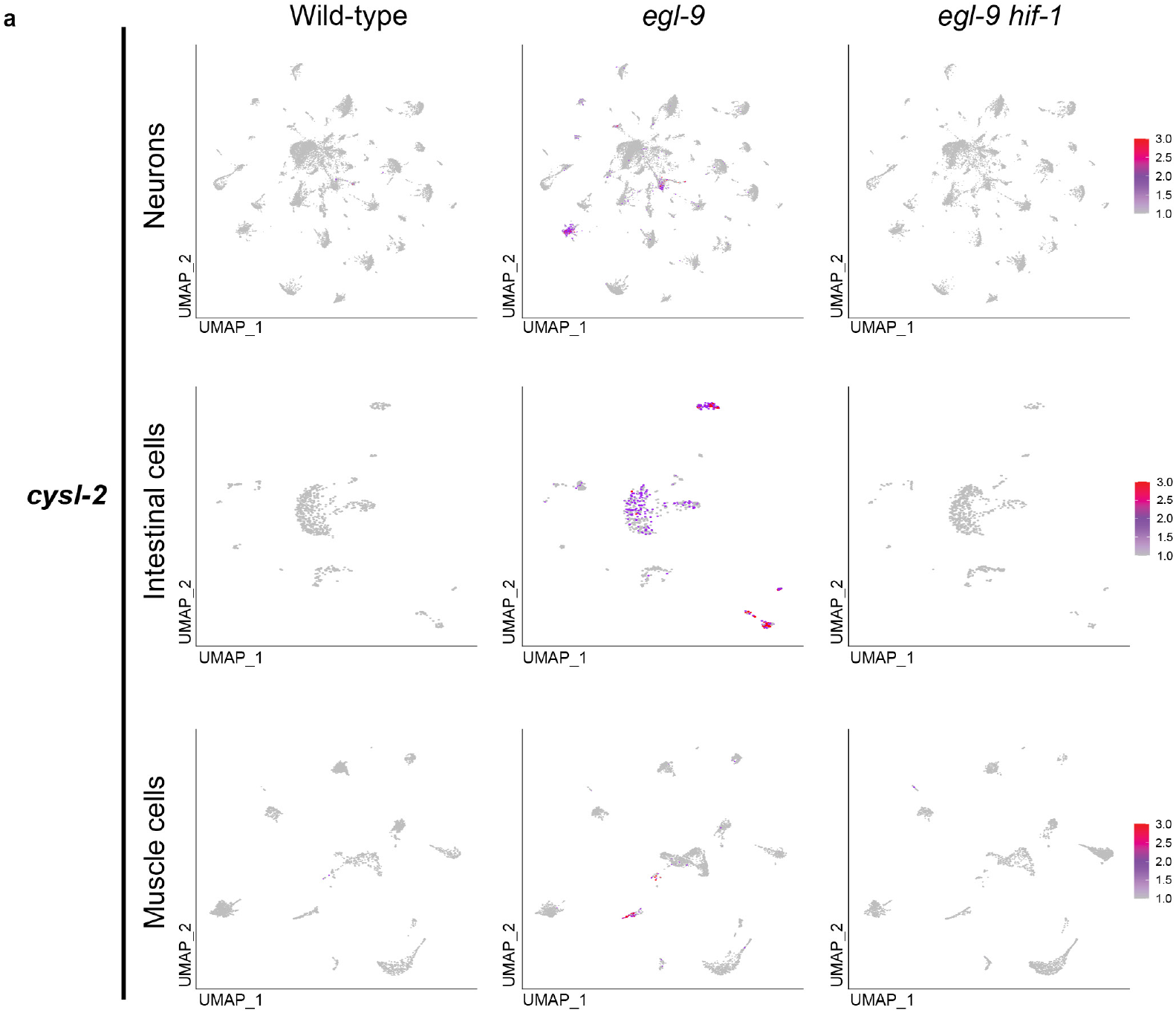
Visualization of *cysl-2* expression in UMAPs of neuronal, intestinal, and muscle tissue by genotype. **a**, UMAPs of neuronal, intestinal, and muscle tissue separated by genotype showing differential expression of *cysl-2* – a gene known to be activated by HIF-1^21^ – in wild-type, *egl-9* mutant, and *egl-9 hif-1* double-mutant animals. Each projection shows the color-coded expression levels of *cysl-2* in cells from that genotype. As expected, *egl-9* mutants exhibited significantly increased expression of *cysl-2* in numerous cell types compared to wild-type or *egl-9 hif-1* double-mutant animals.

**Extended Data Figure 6.**
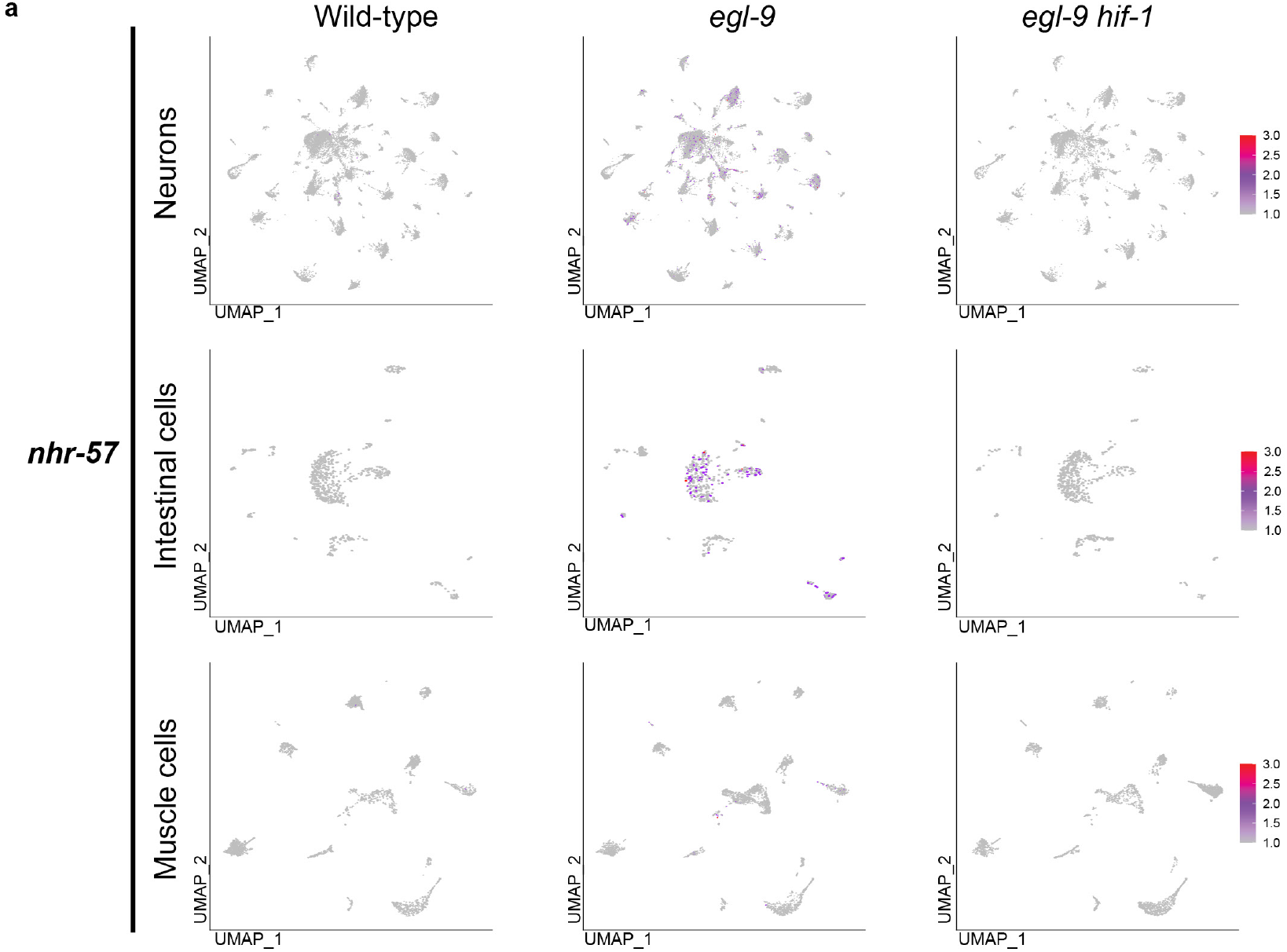
Visualization of *nhr-57* expression in UMAPs of neuronal, intestinal, and muscle tissue by genotype. **a**, UMAPs of neuronal, intestinal, and muscle tissue separated by genotype showing differential expression of *nhr-57* – a gene known to be activated by HIF-1^23^ – in wild-type, *egl-9* mutant, and *egl-9 hif-1* double-mutant animals. Each projection shows the color-coded expression levels of *nhr-57* in cells from that genotype. As expected, *egl-9* mutants exhibited significantly increased expression of *nhr-57* in numerous cell types compared to wild-type or *egl-9 hif-1* double-mutant animals.

**Extended Data Figure 7.**
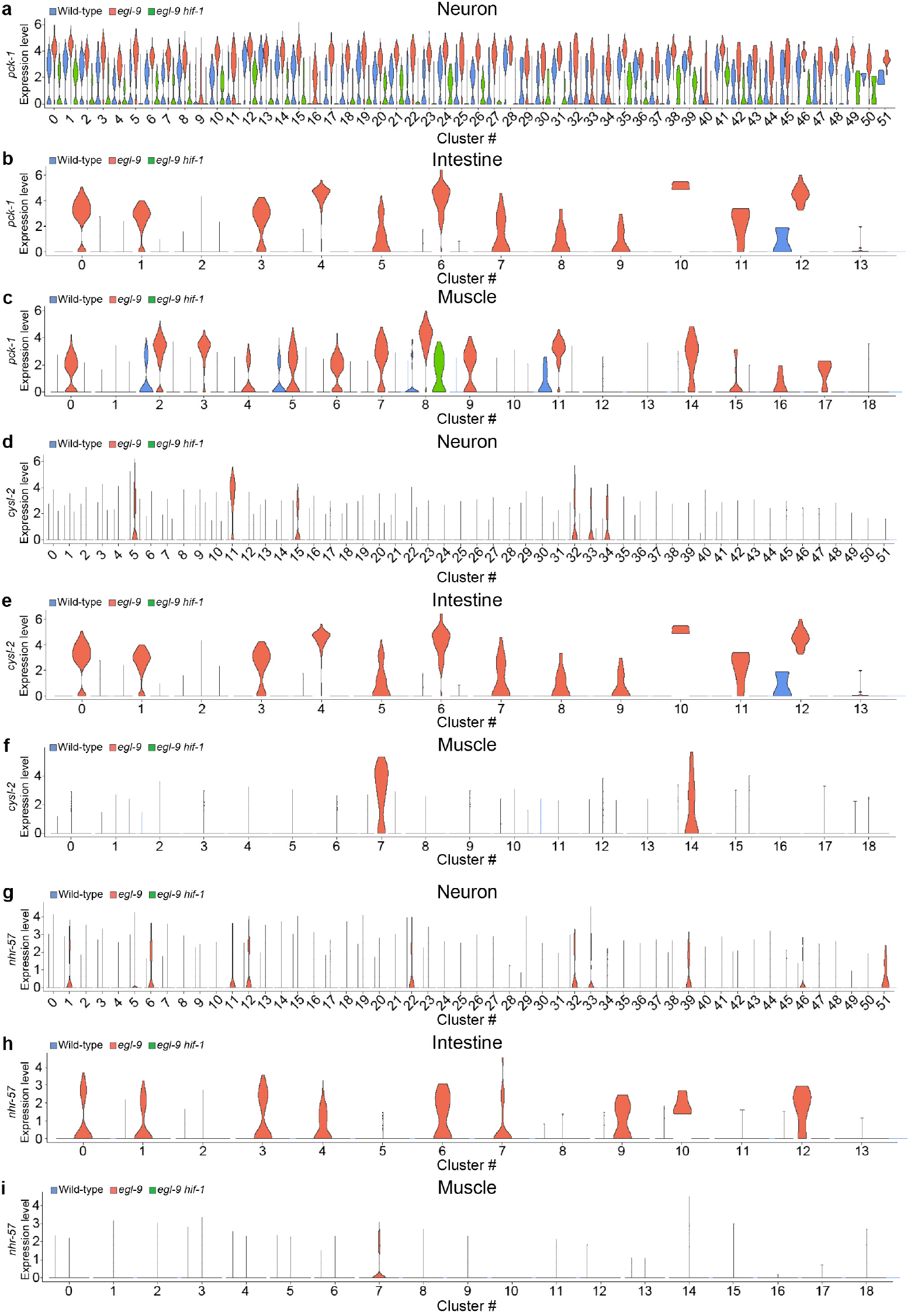
Visualization of *pck-1, cysl-2* and *nhr-57* expression in violin plots of neuronal, intestinal, and muscle tissue by genotype. **a-i**, Violin plots of neuronal, intestinal, and muscle tissue separated by genotype showing differential expression of (**a**-**c**) *pck-*1, (**d**-**f**) *cysl-2* and (**g**-**i**) *nhr-57* – genes known to be activated by HIF-1^21–23^ – in wild-type, *egl-9* mutant, and *egl-9 hif-1* double-mutant animals. Each projection shows the color-coded expression levels of *pck-1, cysl-2* or *nhr-57* in clusters from the indicated genotype. As expected, *egl-9* mutants exhibited significantly increased expression of *pck-1, cysl-2* and *nhr-57* compared to wild-type or *egl-9 hif-1* double-mutant animals, although only certain clusters displayed such HIF-1-dependent regulation.

**Extended Data Figure 8.**
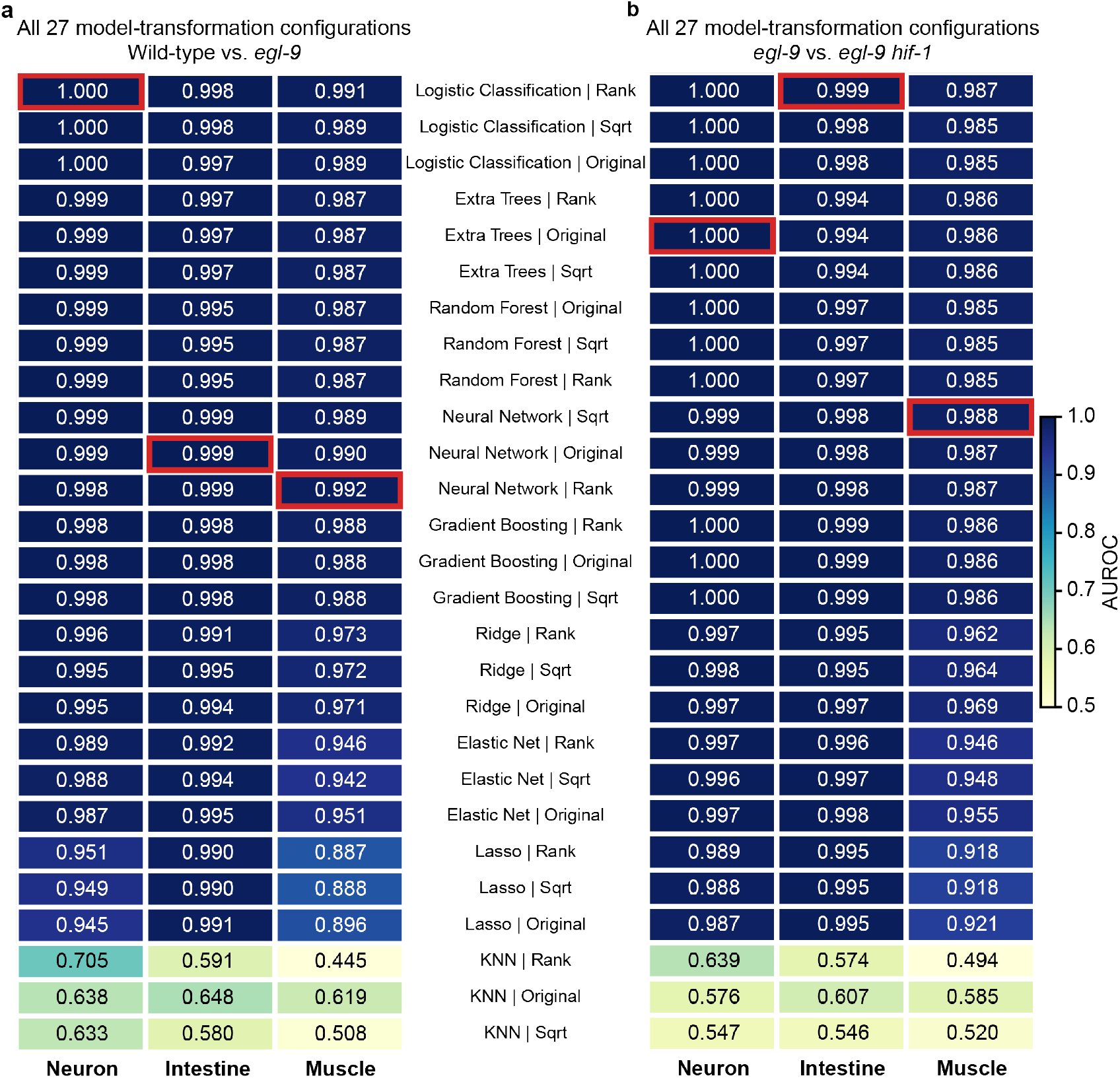
Benchmarking of model–transformation configurations. **a,b**, Mean cross-validated AUROC for all 27 configurations (nine classifiers × three data representations) in neurons, intestinal cells, and muscle cells, for wild type vs. *egl-9* (a) and *egl-9* vs. *egl-9 hif-1* (b). Rows are identical in both panels and ordered by neuronal AUROC in a. Red outlines indicate the model–transformation configurations selected for permutation-importance analysis on the basis of mean five-fold cross-validated AUROC; where configurations were tied, one of the tied configurations was selected (Extended Data Table 3). Values are means across the five cross-validation folds; fold-level standard deviations are provided in Extended Data Table 3.

**Extended Data Figure 9.**
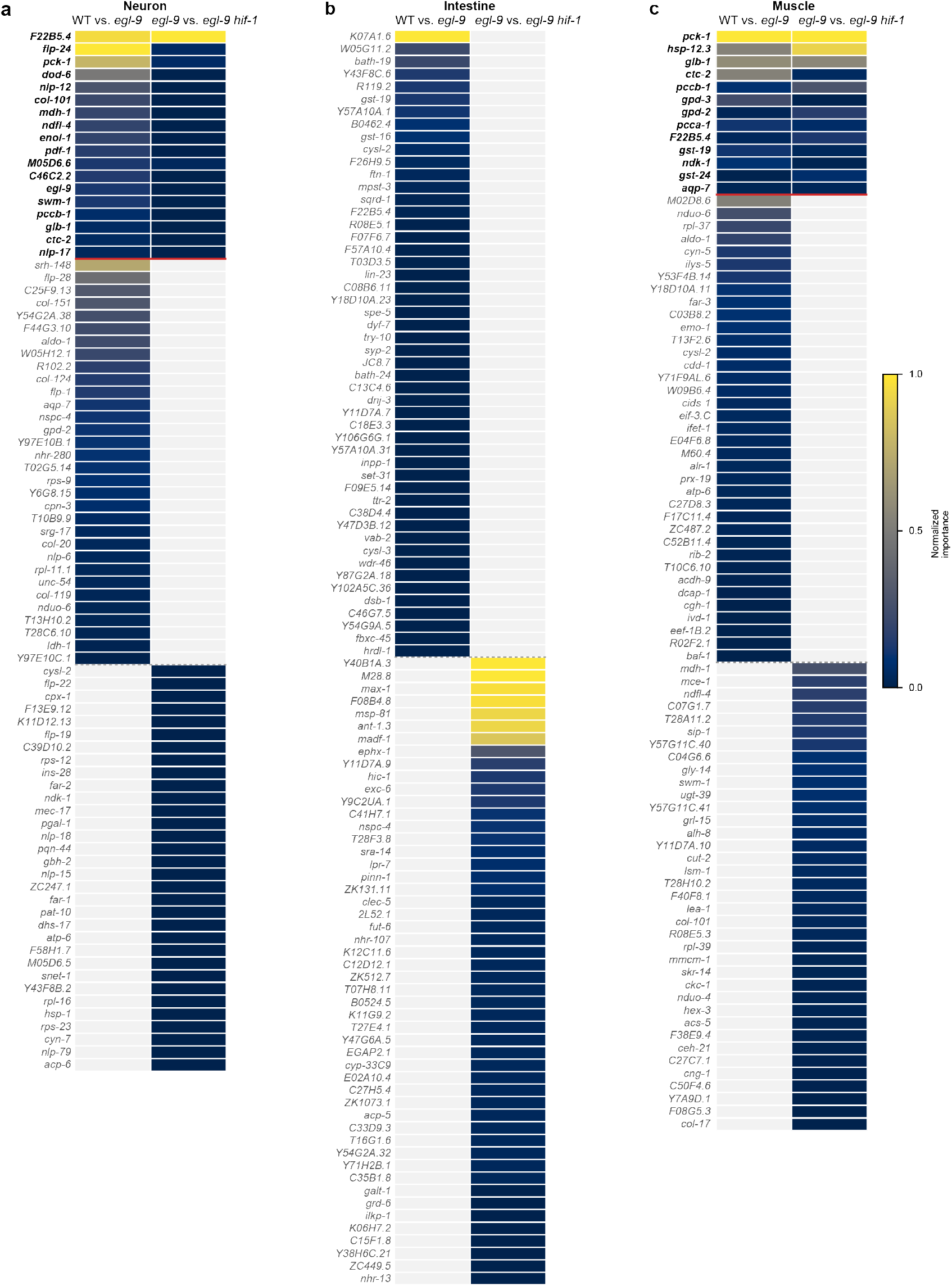
Top predictive genes compared between the two complementary genotype comparisons. **a–c**, The 50 most predictive genes (permutation importance, selected model–transformation configuration per tissue and comparison; Extended Data Table 3) for wild type vs. *egl-9* and *egl-9* vs. *egl-9 hif-1* in neurons (a), intestinal cells (b), and muscle cells (c). Rows show the union of the two top-50 lists: genes identified in both comparisons (bold, above the red line), followed by genes specific to each comparison. Color indicates permutation importance min–max normalized within each comparison’s top 50 (1, most important); gray, gene not among that comparison’s top 50. Shared gene counts: neurons, 18; intestine, 0; muscle, 13. For the intestine, top-gene lists are tie-sensitive near the cutoff, and exact membership should be interpreted with caution.

**Extended Data Figure 10.**
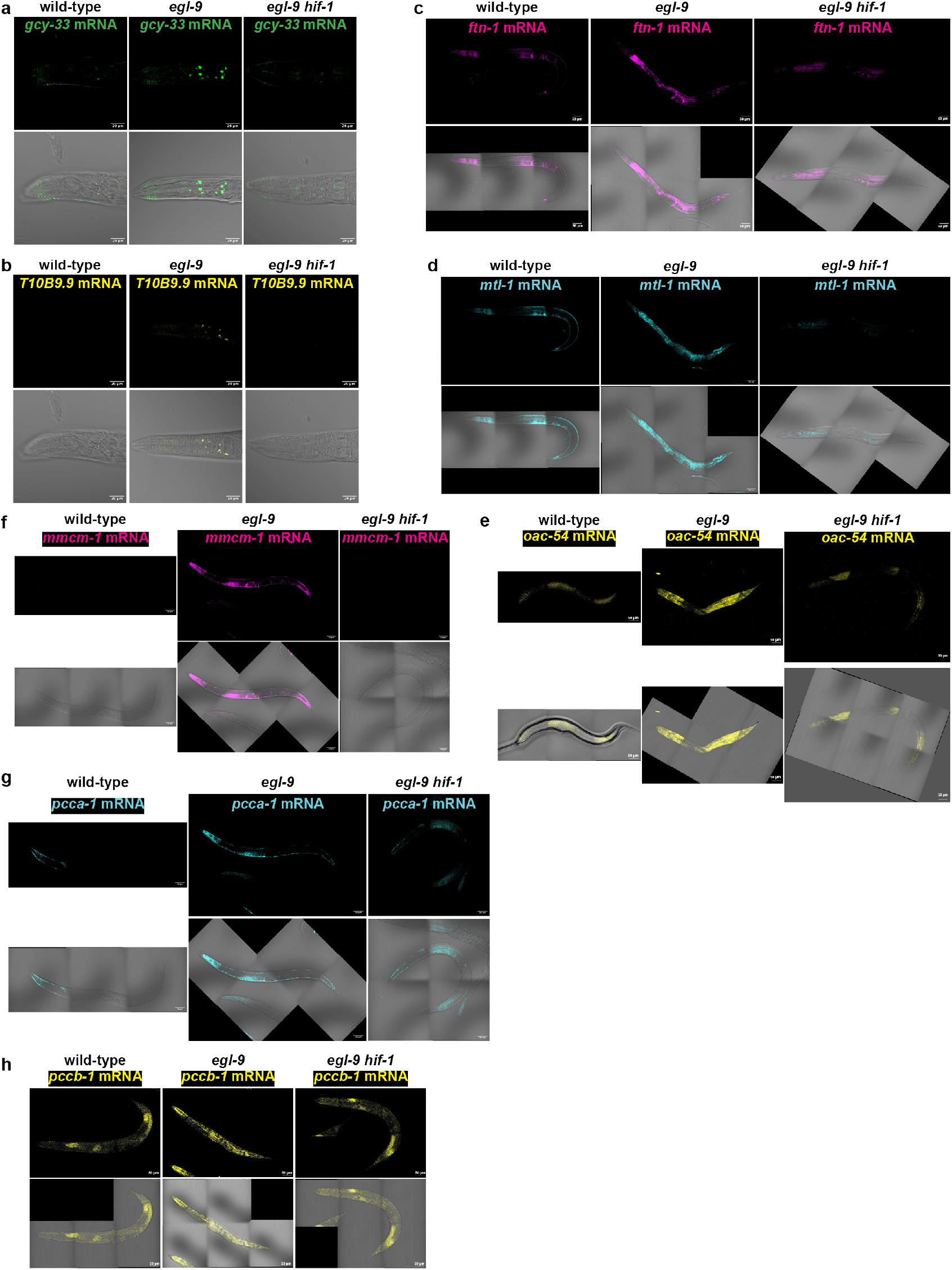
Hybridization Chain Reaction (HCR) RNA fluorescence *in situ* hybridization (FISH) analysis of representative tissue-specific HIF-1 target genes identified by scRNA-Seq. HCR RNA-FISH was used to localize transcripts in young adult wild-type, *egl-9*, and *egl-9 hif-1* animals. Fluorescence images are shown above the corresponding merged brightfield images. **a,b,** In neurons, (a) *gcy-33* and (b) *T10B9.9* mRNA signals were increased in *egl-9* mutants relative to wild-type animals and reduced in *egl-9 hif-1* double mutants. **c–e,** In the intestine, (c) *ftn-1*, (d) *mtl-1*, and (e) *oac-54* mRNA signals were increased in *egl-9* mutants and reduced in *egl-9 hif-1*double mutants. **f–h,** In muscle, (f) *mmcm-1*, (g) *pcca-1*, and (h) *pccb-1* mRNA signals were increased in *egl-9* mutants and reduced in *egl-9 hif-1* double mutants. These expression patterns are consistent with HIF-1-dependent induction in the indicated tissues. Scale bars, 20 µm (a,b) and 50 µm (c–h).

**Extended Data Figure 11.**
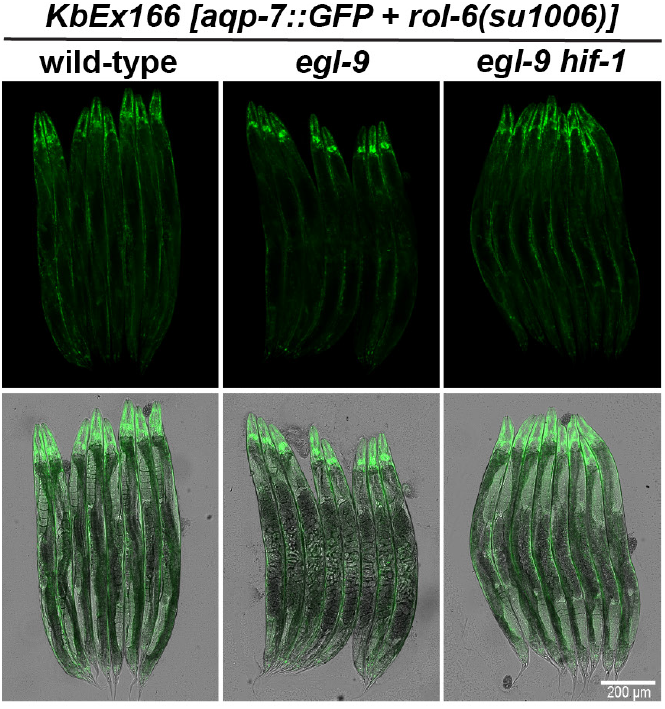
Transcriptional reporter analysis of a neuronal HIF-1-dependent target gene identified by scRNA-Seq. Transcriptional reporter images for *aqp-7*, selected as a representative neuron-enriched HIF-1-dependent DEG from the scRNA-Seq dataset, in day-1 adult wild-type, *egl-9*, and *egl-9 hif-1* animals. Reporter expression was elevated in *egl-9* mutants and reduced in *egl-9 hif-1* double mutants, indicating HIF-1–dependent transcriptional activation. Fluorescence and merged brightfield images are shown. Scale bars, 200 µm. Consistent with the scRNA-Seq data, reporter expression was strongest in neuronal tissues.

**Extended Data Figure 12.**
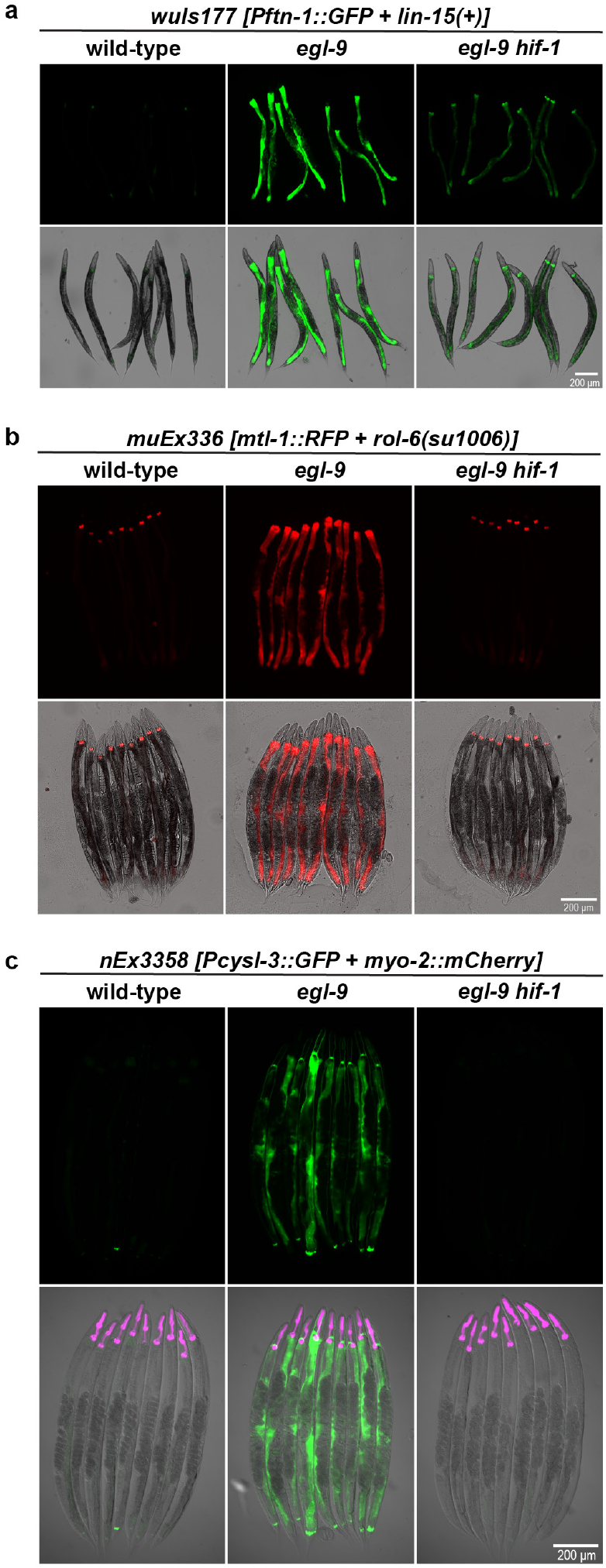
Transcriptional reporter analysis of intestinal HIF-1-dependent target genes identified by scRNA-Seq. **a-c**, Transcriptional reporters for (a) *ftn-1*, (b) *mtl-1* and (c) *cysl-3*, selected as representative intestine-enriched HIF-1-dependent DEGs, imaged in day 1 adult wild-type, *egl-9*, and *egl-9 hif-1* animals. Reporter expression was elevated in *egl-9* relative to wild-type and reduced in *egl-9 hif-1*, consistent with HIF-1–dependent transcriptional activation. Fluorescence (top) and merged brightfield (bottom) images are shown; strain genotypes are indicated above each panel. Scale bars, 200 µm.

**Extended Data Figure 13.**
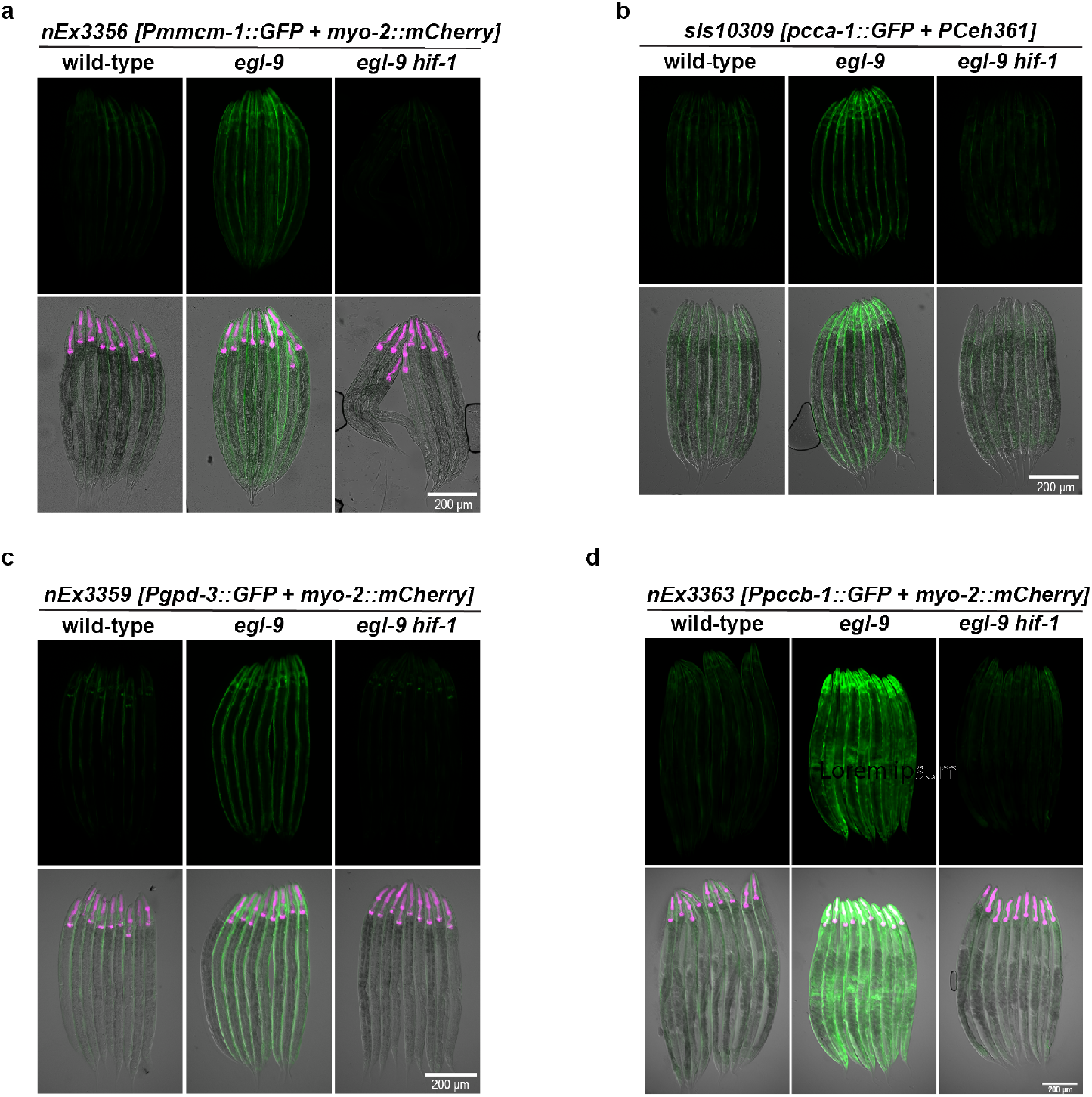
Transcriptional reporter analysis of muscle HIF-1-dependent target genes identified by scRNA-Seq. **a-d**, Transcriptional reporter images for (a) *mmcm-1*, (b) *pcca-1*, (c) *gpd-3*, and (d) *pccb-1*, selected as representative muscle-enriched HIF-1-dependent DEGs from the scRNA-Seq dataset. Reporters in (a-c) were imaged in L4 larvae and (d) in day 1 adults, in wild-type, *egl-9*, and *egl-9 hif-1* animals. Reporter expression was elevated in *egl-9* relative to wild-type and reduced in *egl-9 hif-1*, consistent with HIF-1–dependent transcriptional activation. Fluorescence (top) and merged brightfield (bottom) images are shown; strain genotypes are indicated above each panel. Scale bars, 200 µm.

**Extended Data Figure 14.**
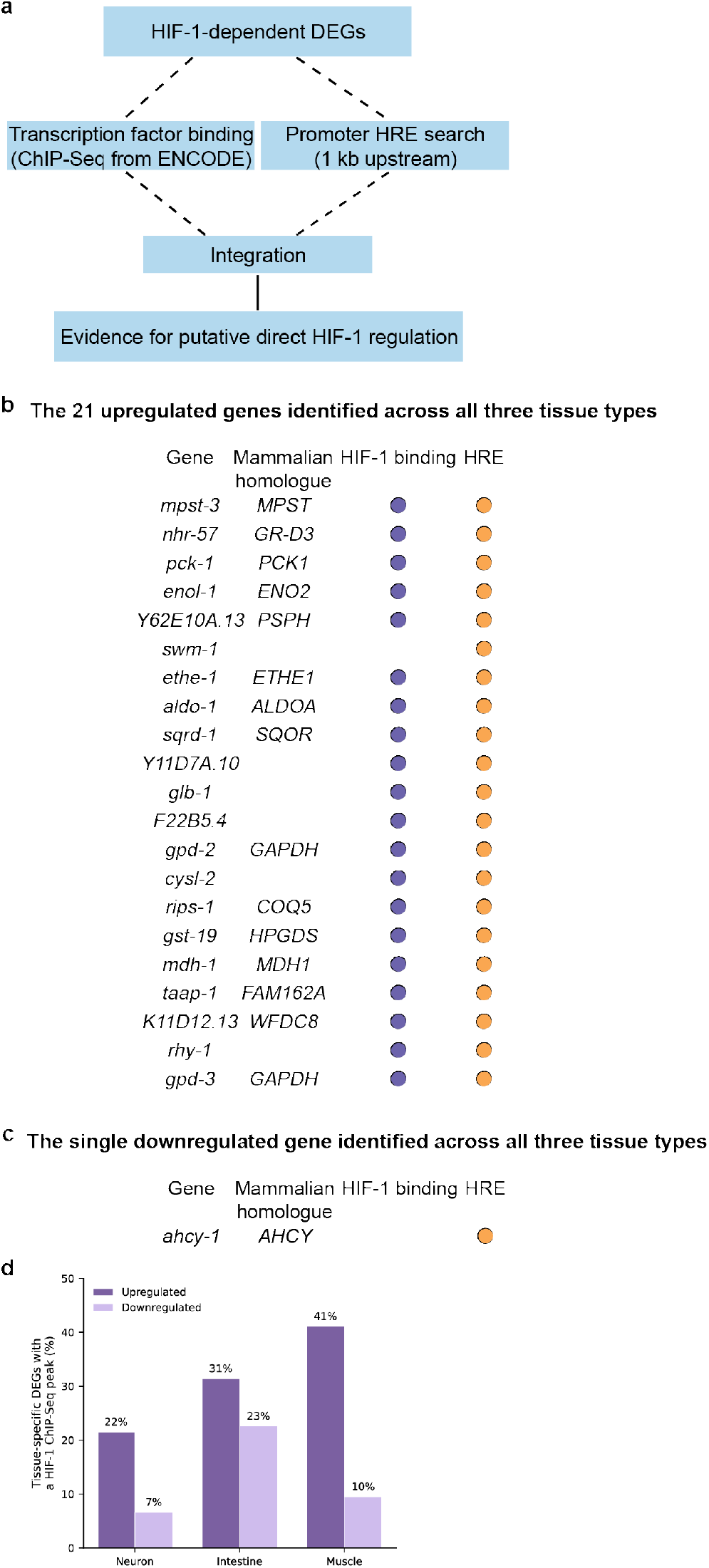
HIF-1 ChIP-Seq binding and promoter HRE analyses identify genes with evidence consistent with direct HIF-1 regulation. **a**, Workflow for comparing HIF-1-dependent DEGs identified by scRNA-Seq with genome-wide HIF-1 ChIP-Seq binding and canonical promoter HRE motifs. HIF-1-dependent DEGs were compared with the HIF-1 ChIP-Seq datasets from ENCODE^22,30^ and with canonical HRE motifs (RCGTG) identified within 1 kb upstream of the transcription start site. These analyses were integrated to assess evidence for putative direct HIF-1 regulation. **b,** List of the 21 genes upregulated in a HIF-1-dependent manner across all three tissues. Corresponding mammalian homologs are listed when available. Dots indicate an associated HIF-1 ChIP-Seq peak or the presence of a promoter-proximal HRE. Twenty of the 21 genes have both HIF-1 ChIP-Seq binding evidence and a promoter HRE; *swm-1* has an HRE but no associated ChIP-Seq peak. **c**, The single gene, *ahcy-1*, downregulated in a HIF-1-dependent manner across all three tissues. *ahcy-1* contains a promoter HRE but lacks an associated HIF-1 ChIP-Seq peak. **d**, Percentage of tissue-specific upregulated and downregulated DEGs associated with a HIF-1 ChIP-Seq peak. Numbers of genes analyzed were 297, 51, and 146 upregulated genes and 350, 31, and 42 downregulated genes for neurons, intestine, and muscle, respectively. Corresponding promoter HRE frequencies are provided in Extended Data Table 2.

**Extended Data Figure 15.**
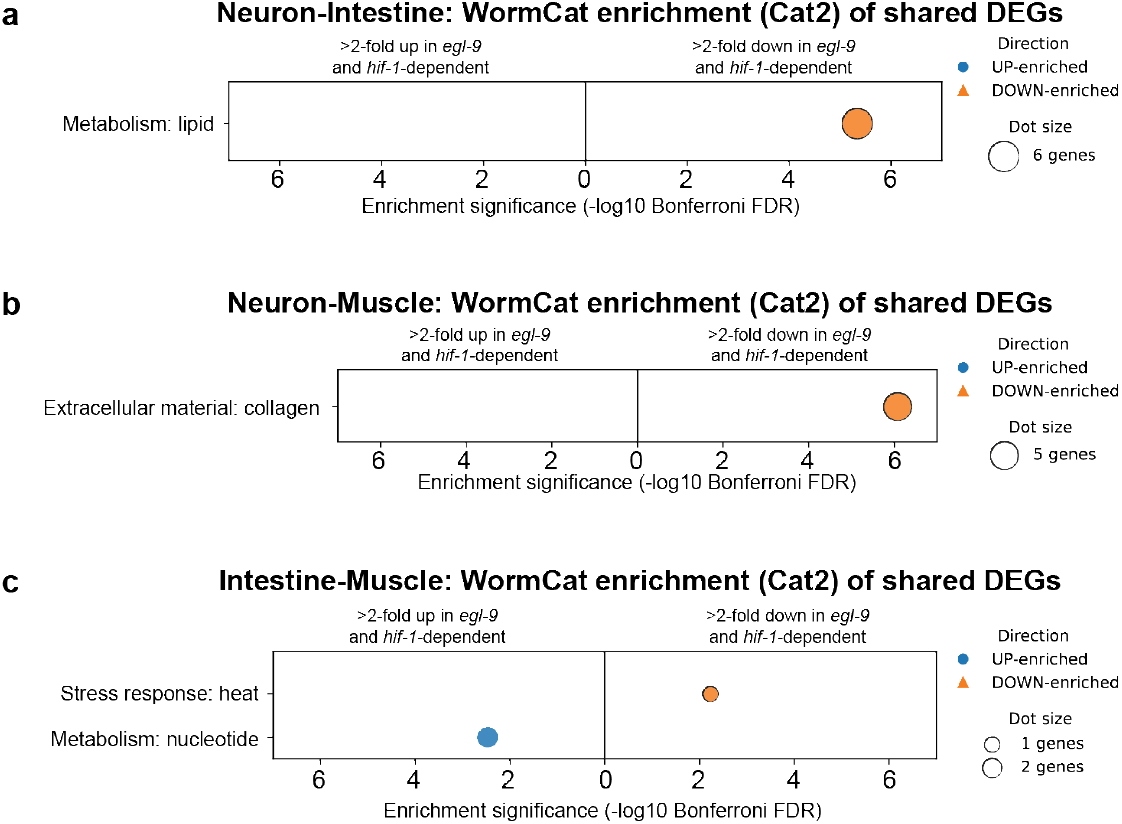
Shared HIF-1-dependent transcriptional responses between tissue pairs are confined to discrete functional categories. **a-c**, WormCat Category 2 (Cat2) enrichment analysis of genes shared between pairs of tissues following HIF-1 activation. Only Bonferroni-significant categories (Bonferroni ≤ 0.05) are shown. For each tissue pair, enriched pathways are displayed separately for shared upregulated (left panels) and shared downregulated (right panels) gene sets. Dot position indicates enrichment significance (−log10 Bonferroni), dot size reflects the number of genes in the overlapping set assigned to each category (RGS), and numbers adjacent to dots indicate RGS values. Across tissue pairs, shared transcriptional responses are limited to discrete functional modules, including **(a)** lipid metabolism (neuron– intestine), **(b)** extracellular matrix components (neuron–muscle), and **(c)** nucleotide/stress-associated pathways (intestine–muscle), indicating that overlap between tissues is pathway-restricted rather than broadly homogeneous.

**Extended Data Figure 16.**
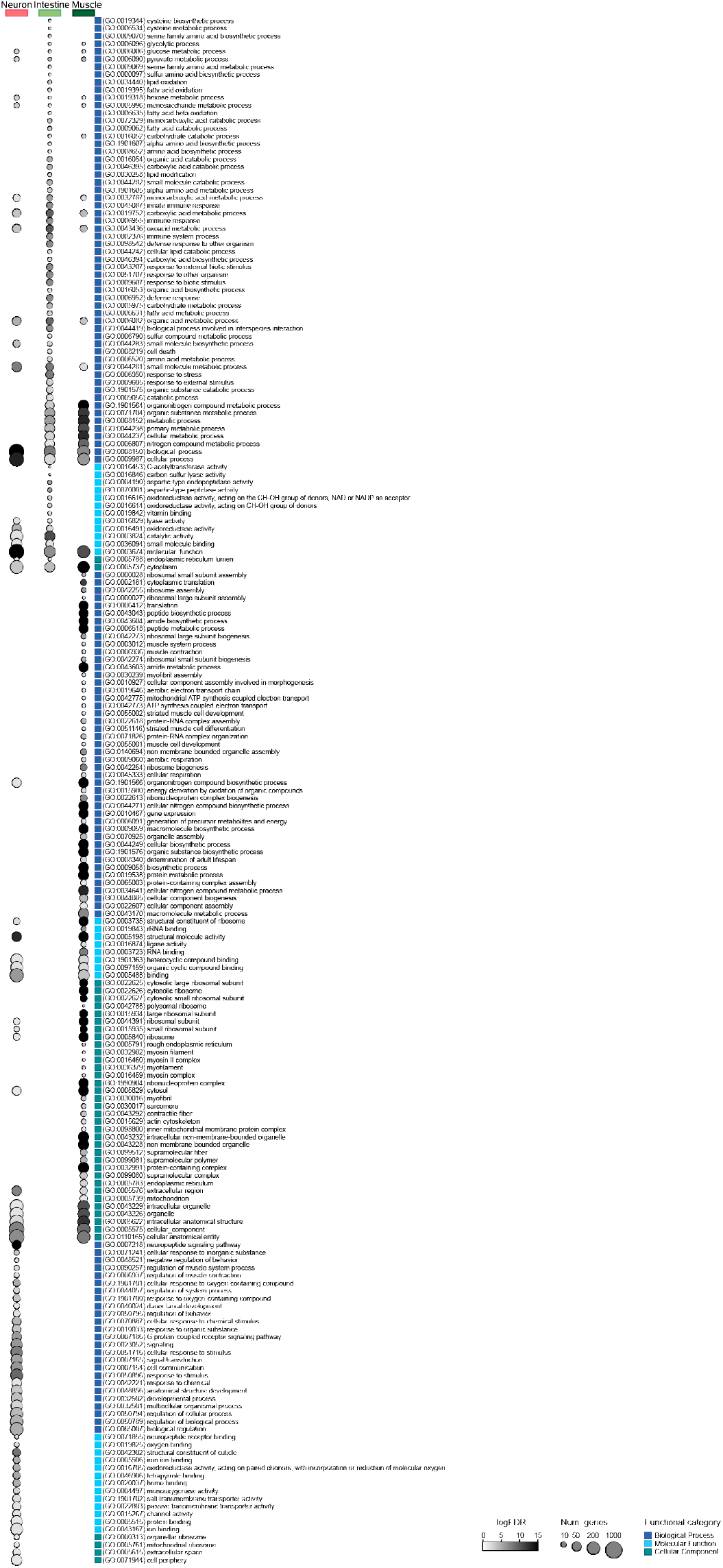
HIF-1 function activates diverse cell biological pathways across cell types. **a**, Gene ontology (GO)-term analyses of HIF-1 activity-dependent genes across the three tissues classified by biological process, molecular function, or cellular component.

**Extended Data Figure 17.**
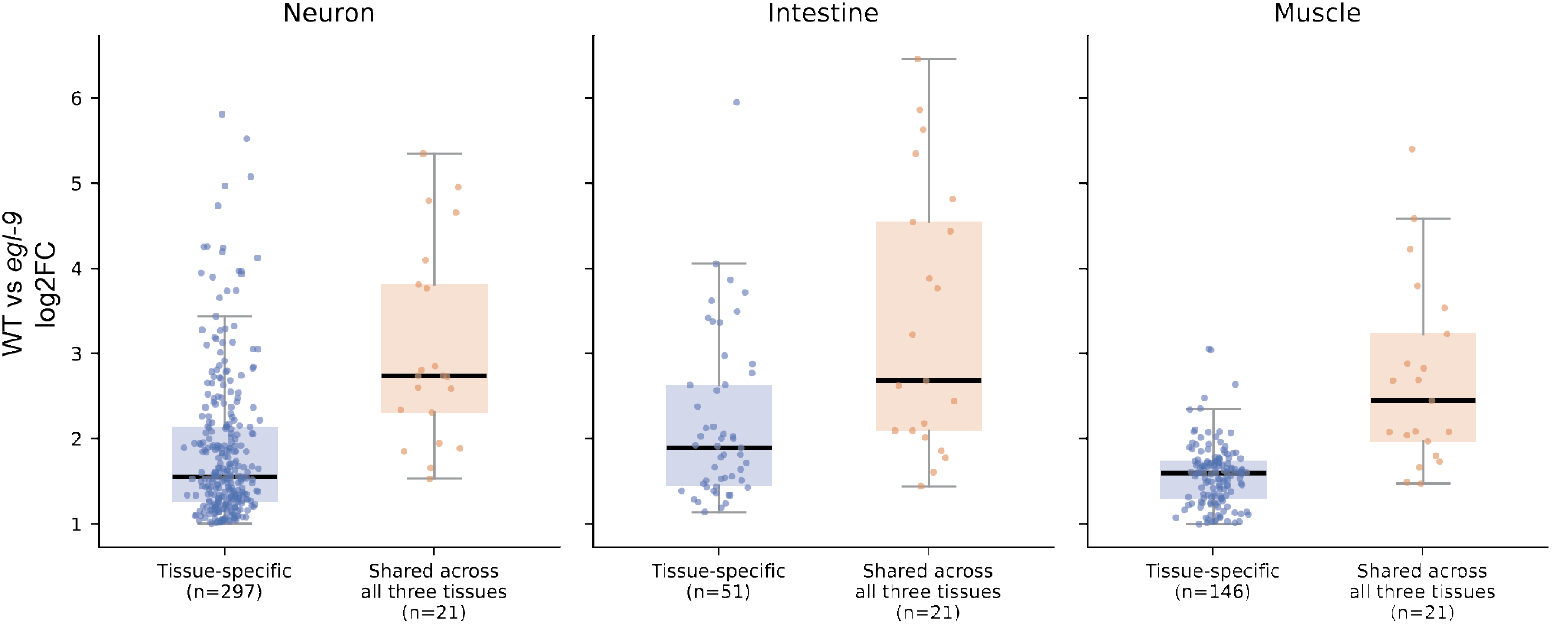
Comparison of tissue-specific and three-tissue-shared HIF-1 responses. HIF-1-dependent induction was compared between genes upregulated exclusively in one tissue and genes upregulated across all three tissues, shown separately for neurons, intestine, and muscle. For each gene, induction was defined as the maximum *egl-9* versus wild-type log2 fold change among clusters meeting the reciprocal criteria for HIF-1 dependence. Each point represents one gene; boxes show the median and interquartile range, and whiskers extend to 1.5 times the interquartile range. The tissue-specific groups contained 297 neuronal, 51 intestinal, and 146 muscle genes; the three-tissue-shared group contained 21 genes. Genes upregulated in two of the three tissues—32 neuronal, 21 intestinal, and 27 muscle genes—were excluded from this comparison. No statistical test was applied; the analysis is descriptive.

**Extended Data Figure 18.**
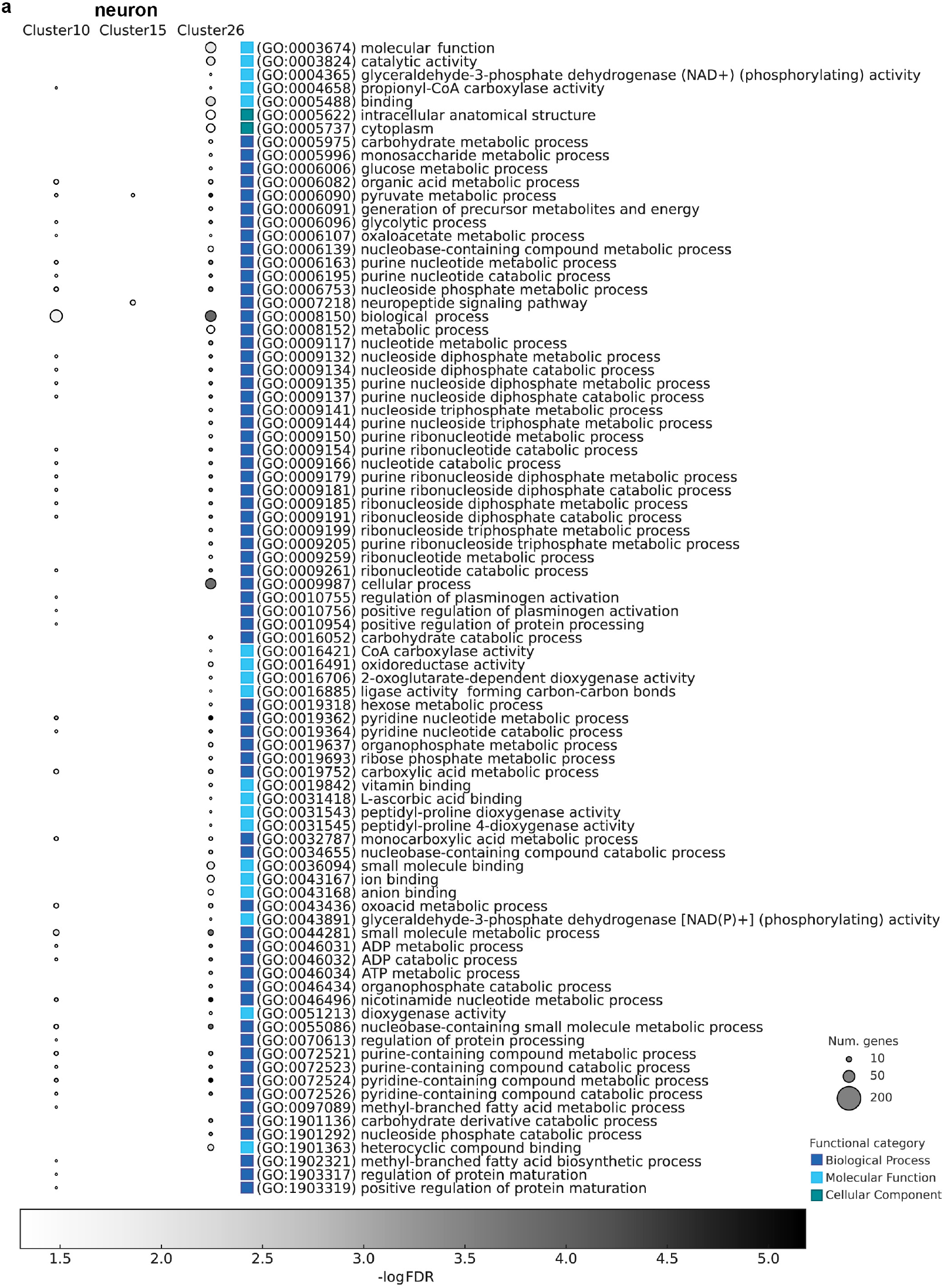
Cluster-specific GO enrichment analysis of HIF-1–dependent genes in neurons. **a**, Gene Ontology (GO) term enrichment analysis of genes that are differentially expressed in a HIF-1–dependent manner across three identified neuronal clusters from scRNA-Seq data. Cluster 15 represents oxygen-sensing BAG neurons, and clusters 10 and 26 represent mechanosensory touch receptor neurons. These clusters were selected for comparison because they represent functionally distinct neuronal classes with sufficient numbers of HIF-1-dependent DEGs for enrichment analysis. GO terms are grouped by functional category: biological process (blue), molecular function (cyan), and cellular component (green).

**Extended Data Figure 19.**
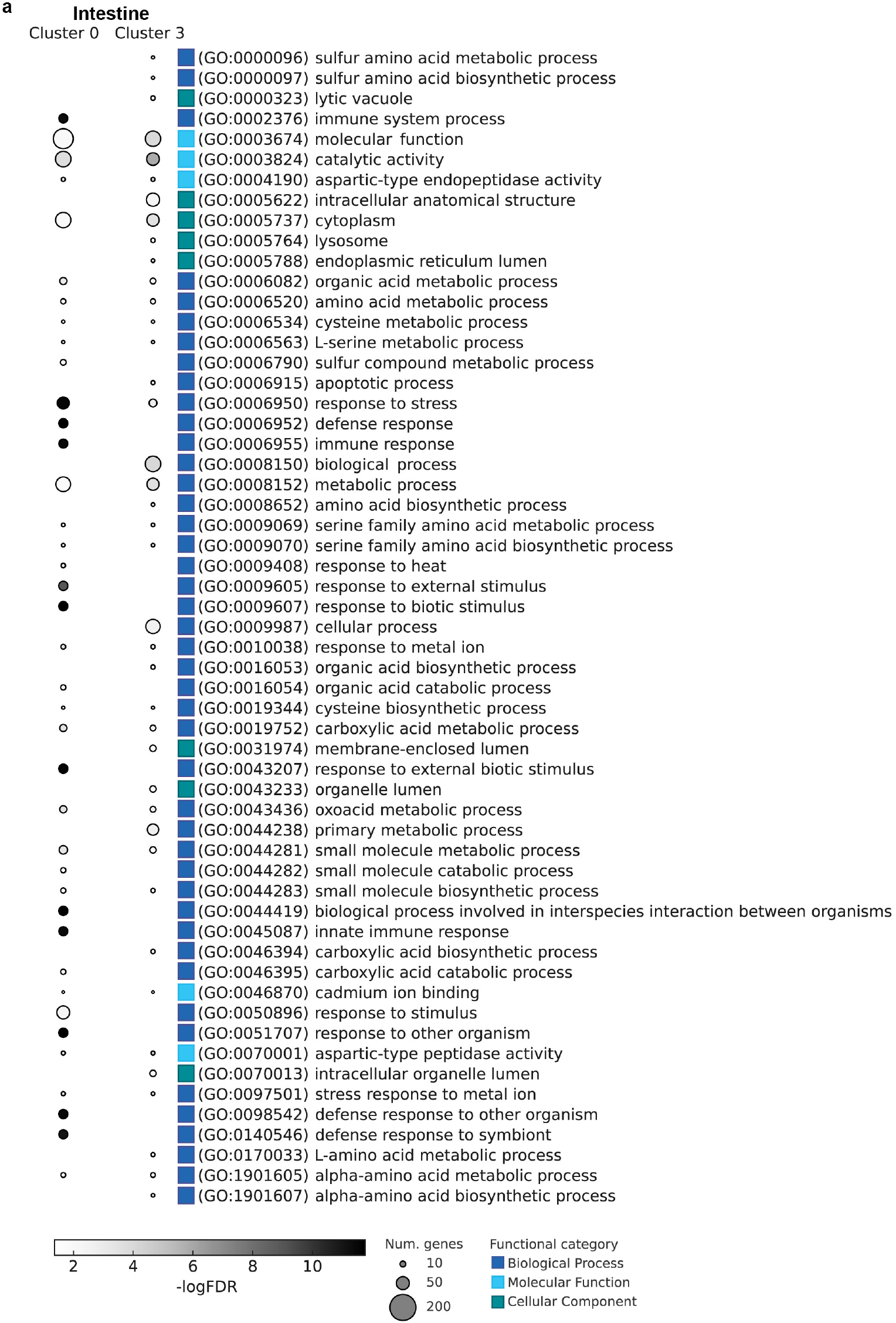
Cluster-specific GO enrichment analysis of HIF-1–dependent genes in intestine. **a**, Gene Ontology (GO) term enrichment analysis of genes that are differentially expressed in a HIF-1–dependent manner across two intestine clusters from scRNA-Seq data. In Extended Data Fig. 3a,b, clusters 0 and 3 were identified as intestinal cell populations based on expression of intestinal markers, *vha-6* and *asp-1*, and Extended Data Fig. 21 further suggests that cluster 3 corresponds predominantly to posterior intestinal cells based on enrichment of the posterior intestinal marker *nob-1*. GO terms are grouped by functional category: biological process (blue), molecular function (cyan), and cellular component (green).

**Extended Data Figure 20.**
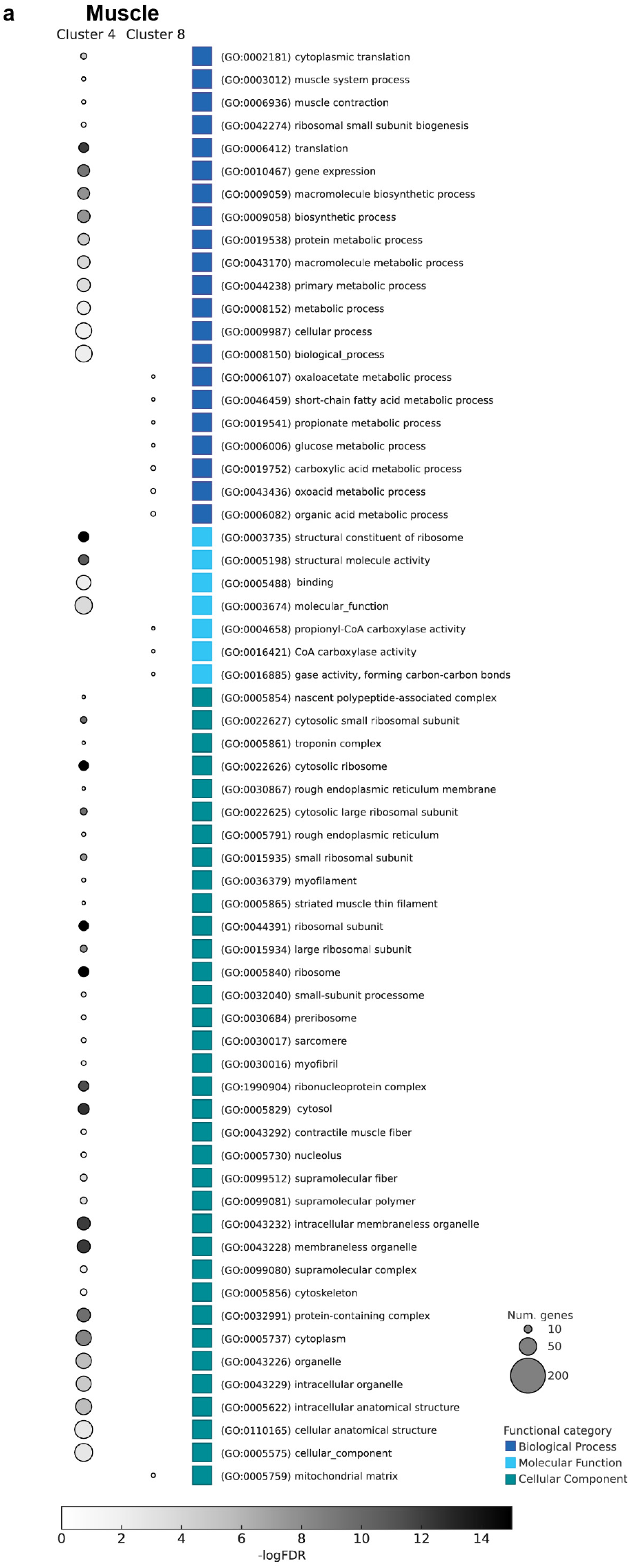
Cluster-specific GO enrichment analysis of HIF-1–dependent genes in muscle. **a**, Gene Ontology (GO) term enrichment analysis of genes that are differentially expressed in a HIF-1–dependent manner across two muscle clusters from scRNA-Seq data. Cluster 4 represents vulval muscle cells, and cluster 8 represents anterior body wall muscle cells (see text). GO terms are grouped by functional category: biological process (blue), molecular function (cyan), and cellular component (green).

**Extended Data Figure 21.**
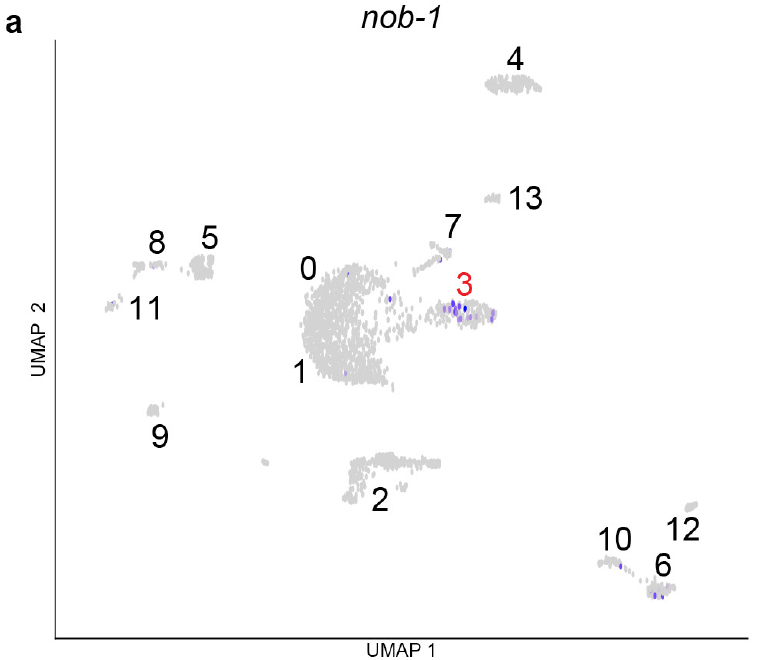
Cluster 3 likely represents posterior intestinal cells. **a**, UMAP projection of intestine cells identified using posterior intestine cell-specific marker *nob-1* highlighted in purple^46,51^. In Extended Data Fig. 3a,b, clusters 0 and 3 were identified as intestinal cell clusters based on expression of intestinal markers. Enrichment of *nob-1*-expressing cells within Cluster 3 further suggests that this cluster represents predominantly posterior intestinal cells.

**Extended Data Figure 22.**
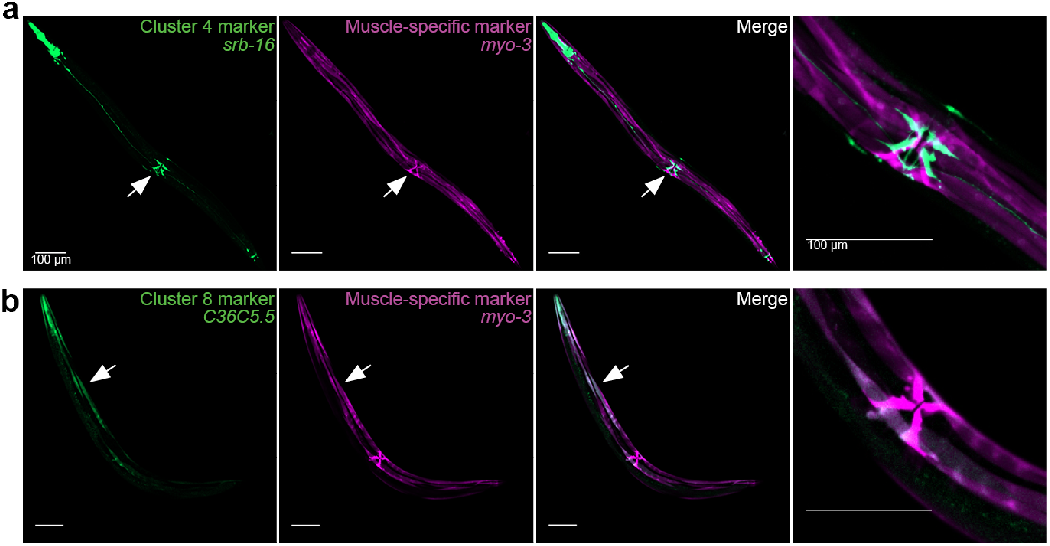
Spatial expression of muscle cluster 4 marker gene and cluster 8 marker genes. **a-b**, GFP expression, (**a**) driven by the *srb-16* promoter (cluster 4 marker gene) and (**b**) the *C36C5.5* promoter (cluster 8 marker gene), is overlaid with mCherry expression driven by the muscle-specific *myo-3* promoter, which was also used to sort muscle cells. This overlay identified muscle clusters 4 and 8 as likely corresponding to the vulval muscles and anterior body-wall muscles, respectively, indicated by white arrows. More specifically, to determine the identity of the cells represented by muscle cluster 4, we examined our atlas for genes that might serve as anatomical markers based on their cluster-4-specific expression and identified the gene *srb-16* as described in Extended Data Fig. 22a. We generated a GFP transcriptional reporter for *srb-16* (*P_srb-16_::gfp)* and compared its expression pattern with that of an mCherry reporter driven by the promoter for the muscle-cell marker *myo-3* (*P_myo-3_::mCherry*), which we had used to sort muscle cells. The *srb-16* reporter labeled vulval and pharyngeal muscles as well as some unidentified neurons and overlapped most strikingly with the *myo-3* marker in the worm’s eight vulval muscle cells (Extended Data Fig. 22a). We used an analogous approach to determine the identity of muscle cluster 8. A muscle cluster 8 marker gene *C36C5.5* reporter overlapped with the *myo-3* marker in anterior but not posterior body-wall muscle cells (Extended Data Fig. 22b), revealing that different muscle UMAP clusters can represent anatomically distinct cell subsets within an apparently homogeneous muscle tissue. In short, based on marker gene expression patterns, cluster 4 represents vulval muscles, while cluster 8 represents anterior body-wall muscles.

**Extended Data Figure 23.**
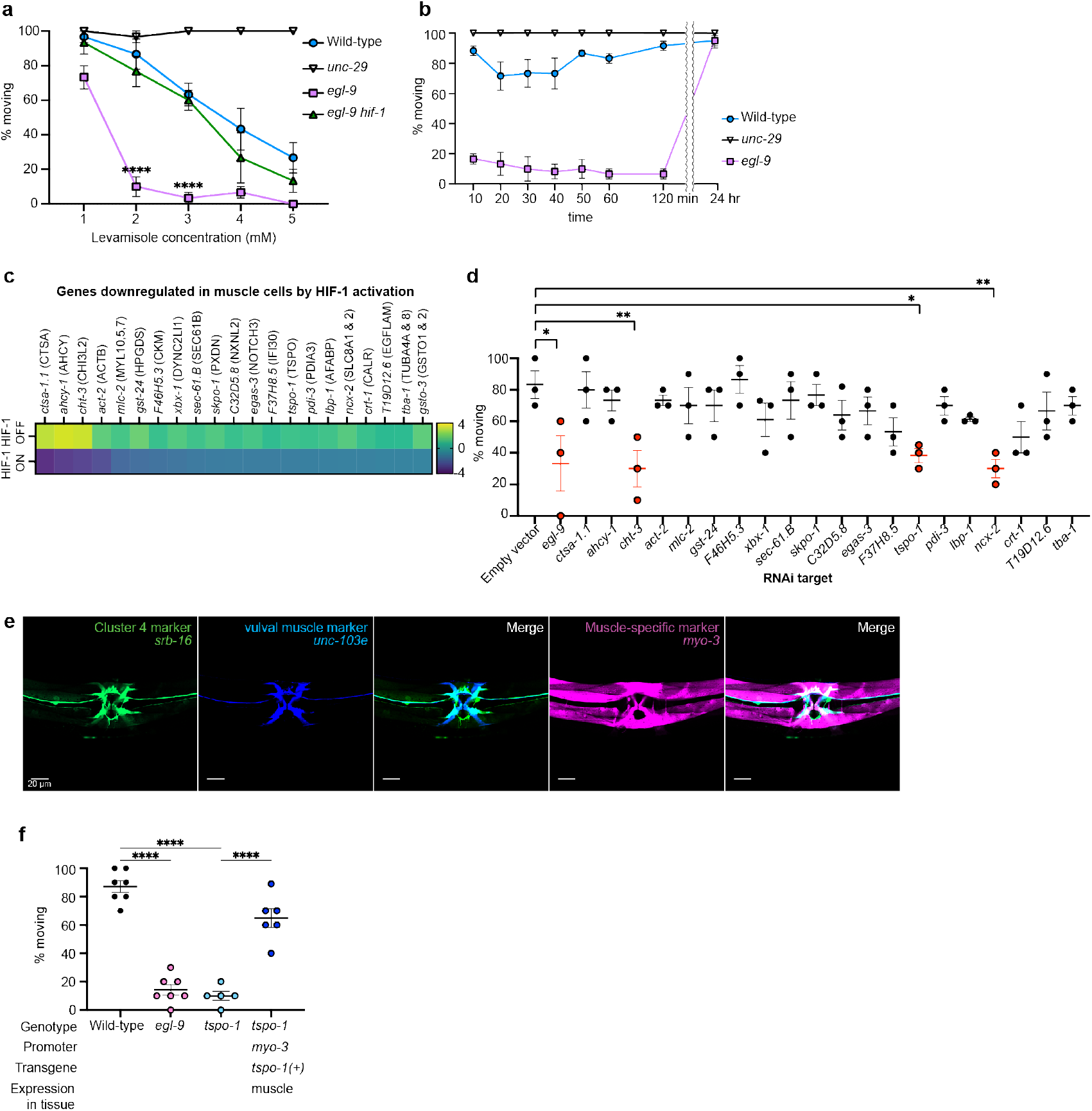
Functional identification of *tspo-1* as a subtype-specific effector of HIF-1-dependent muscle physiology. **a**, Dose-dependent levamisole sensitivity.Approximately 10 adult worms were exposed to the indicated concentration of levamisole, and the percentage of animals moving after 10 min was quantified. *unc-29* encodes a subunit of the levamisole-sensitive nicotinic acetylcholine receptor and *unc-29* mutants are levamisole-insensitive^39^. *egl-9* mutants exhibited hypersensitivity to levamisole compared to wild-type animals, and this hypersensitivity required *hif-1(+)* function. Data points and error bars represent the mean ± s.e.m. from at least three independent assays. **b,** Levamisole equilibrium assay. Approximately 10 adult worms were exposed to a 2 mM levamisole drop, and the percentage of animals moving was quantified at the indicated time points following exposure. The levamisole drop had dried before the 10-min time point. *egl-9* mutants exhibited pronounced hypersensitivity to levamisole, with substantially reduced mobility that persisted through 120 min after exposure; movement was largely restored by 24 hr, indicating that the *egl-9* hypersensitive phenotype is reversible. Wild-type animals retained substantial mobility throughout the time course, whereas *unc-29* mutants remained largely insensitive to levamisole. Data are presented as mean ± s.e.m. **c,** Heatmap showing evolutionarily conserved genes downregulated in muscle cells by HIF-1 activation. Genes are ordered according to the degree of HIF-1-dependent downregulation observed in *egl-9* mutant muscle cells. The “HIF-1 ON” row indicates the log2 fold change in expression between wild-type and *egl-9* mutant muscle cells (wild-type / *egl-9*), whereas the “HIF-1 OFF” row indicates the log2 fold change between *egl-9* mutant and *egl-9 hif-1* double-mutant muscle cells (*egl-9* / *egl-9 hif-1*). Color scale indicates log2 fold change in expression level. **d,** Muscle-specific RNAi^45^ screen for modulators of levamisole sensitivity. A strain in which RNAi activity was restricted to muscle tissue was fed bacteria expressing RNAi constructs targeting candidate genes identified from panel c. Approximately 10 adult worms were exposed to 2 mM levamisole, and the percentage of animals moving after 10 min was quantified. Feeding bacteria carrying an empty vector control did not alter levamisole sensitivity compared to wild-type animals. Muscle-specific knockdown of *egl-9* elicited hypersensitivity to levamisole similar to that observed in *egl-9* mutants. Knockdown of *cht-3*, *tspo-1*, or *ncx-2* also resulted in pronounced levamisole hypersensitivity and these genes were prioritized for further analysis. Data points represent independent assays, and error bars indicate mean ± s.e.m. Data points and error bars for *egl-9*, *cht-3*, *tspo-1*, and *ncx-2* are shown in red. **e,** GFP expression driven by the cluster 4 marker *srb-16* overlapped with BFP expression from the vulval-muscle-specific promoter *unc-103e (Punc-103e::bfp)*^52^ and mCherry expression driven by the muscle-specific promoter *myo-3*^17^. The overlap in expression patterns from the *srb-16*, *unc-103e*, and *myo-3* promoters indicates that cluster 4 corresponds to vulval muscle cells. **f,** *egl-9* and *tspo-1* null mutants are hypersensitive to levamisole. Approximately 10 adult worms were exposed to 2 mM levamisole, and the percentage of animals moving after 10 min was quantified. Expression of *tspo-1(+)* in muscle tissue using the *myo-3* promoter rescued the levamisole hypersensitivity of *tspo-1* null-mutant animals.

**Extended Data Figure 24.**
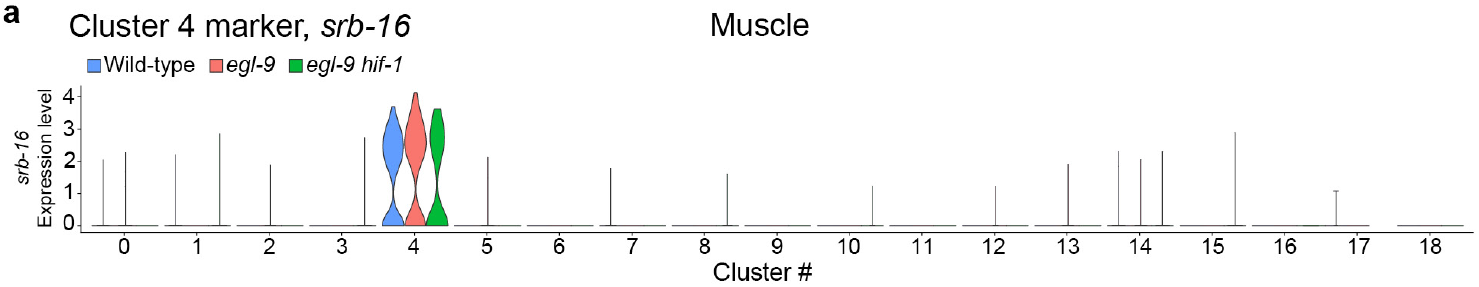
Expression of putative cluster 4 marker *srb-16* in muscle tissue. **a**, Violin plot depicting *srb-16* expression levels across all clusters from muscle tissue from wild-type, *egl-9* mutant, and *egl-9 hif-1* double-mutant animals. Note that *srb-16* appears to be robustly expressed only within muscle cluster 4 and across all genotypes.

**Extended Data Figure 25.**
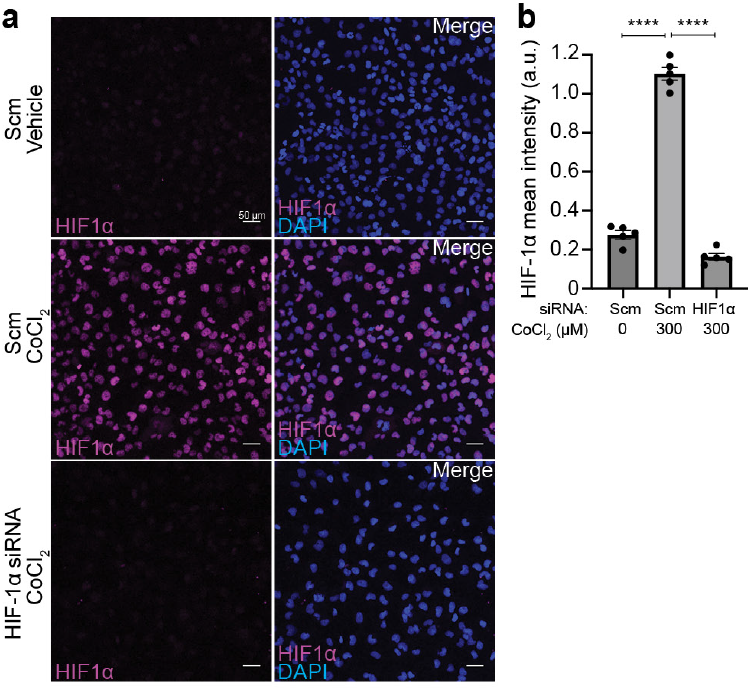
siRNA against HIF-1α reduces cobalt chloride-induced HIF-1α levels in human AC16 cardiomyocytes. **a**, Immunocytochemistry of HIF-1α protein levels in AC16 cardiomyocytes treated with vehicle or 300 µM cobalt chloride (CoCl2) in the presence of scrambled control siRNA (Scm) or siRNA targeting HIF-1α. DAPI was used to label nuclei. **b,** Quantification of HIF-1α immunofluorescence intensity shown in panel a (*n* = 5 images per condition). Treatment with CoCl2 increased HIF-1α levels in cells treated with scrambled control siRNA, whereas siRNA targeting HIF-1α markedly reduced CoCl2-induced HIF-1α levels. Mean fluorescence intensity was quantified per image and normalized to DAPI fluorescence intensity to account for nuclear content. Data are presented as mean ± s.e.m.

## Extended Data Tables 1 to 6

Table S1. Tissue-specific differential expression

Table S2. Integration of DEGs and HIF-1 binding features

Table S3. ML model benchmarking

Table S4. WormCat enrichment results

Table S5. GO enrichment results

Table S6. Strain list

## References

1. Maxwell, P. H. et al. The tumour suppressor protein VHL targets hypoxia-inducible factors for oxygen-dependent proteolysis. Nature 399, 271–275 (1999).

2. Jaakkola, P. et al. Targeting of HIF-alpha to the von Hippel-Lindau ubiquitylation complex by O2-regulated prolyl hydroxylation. Science 292, 468–472 (2001).

3. Ivan, M. et al. HIFalpha targeted for VHL-mediated destruction by proline hydroxylation: implications for O2 sensing. Science 292, 464–468 (2001).

4. Epstein, A. C. et al. C. elegans EGL-9 and mammalian homologs define a family of dioxygenases that regulate HIF by prolyl hydroxylation. Cell 107, 43–54 (2001).

5. Semenza, G. L. Hypoxia-inducible factors in physiology and medicine. Cell 148, 399–408 (2012).

6. Kaelin, W. G. The VHL Tumor Suppressor Gene: Insights into Oxygen Sensing and Cancer. Trans Am Clin Climatol Assoc 128, 298–307 (2017).

7. Wheaton, W. W. & Chandel, N. S. Hypoxia. 2. Hypoxia regulates cellular metabolism. Am J Physiol Cell Physiol 300, C385–393 (2011).

8. Harris, A. L. Hypoxia--a key regulatory factor in tumour growth. Nat Rev Cancer 2, 38–47 (2002).

9. Bianciardi, P. et al. Chronic in vivo hypoxia in various organs: hypoxia-inducible factor-1alpha and apoptosis. Biochem Biophys Res Commun 342, 875–880 (2006).

10. Stroka, D. M. et al. HIF-1 is expressed in normoxic tissue and displays an organ-specific regulation under systemic hypoxia. FASEB J 15, 2445–2453 (2001).

11. Trent, C., Tsuing, N. & Horvitz, H. R. Egg-laying defective mutants of the nematode *Caenorhabditis elegans*. Genetics 104, 619–647 (1983).

12. Jiang, H., Guo, R. & Powell-Coffman, J. A. The *Caenorhabditis elegans hif-1* gene encodes a bHLH-PAS protein that is required for adaptation to hypoxia. Proceedings of the National Academy of Sciences 98, 7916– 7921 (2001).

13. Sulston, J. E., Schierenberg, E., White, J. G. & Thomson, J. N. The embryonic cell lineage of the nematode *Caenorhabditis elegans*. Dev Biol 100, 64–119 (1983).

14. Sulston, J. E. & Horvitz, H. R. Post-embryonic cell lineages of the nematode, *Caenorhabditis elegans*. Dev Biol 56, 110–156 (1977).

15. Altun-Gultekin, Z. et al. A regulatory cascade of three homeobox genes, *ceh-10*, *ttx-3* and *ceh-23*, controls cell fate specification of a defined interneuron class in *C. elegans*. Development 128, 1951–1969 (2001).

16. Oka, T., Toyomura, T., Honjo, K., Wada, Y. & Futai, M. Four subunit a isoforms of *Caenorhabditis elegans* vacuolar H+-ATPase. Cell-specific expression during development. J Biol Chem 276, 33079–33085 (2001).

17. Fox, R. M. et al. The embryonic muscle transcriptome of *Caenorhabditis elegans*. Genome Biol 8, R188 (2007).

18. Taylor, S. R. et al. Molecular topography of an entire nervous system. Cell 184, 4329–4347.e23 (2021).

19. Tcherepanova, I., Bhattacharyya, L., Rubin, C. S. & Freedman, J. H. Aspartic proteases from the nematode *Caenorhabditis elegans*. Structural organization and developmental and cell-specific expression of *asp-1*. J Biol Chem 275, 26359–26369 (2000).

20. Kharchenko, P. V., Silberstein, L. & Scadden, D. T. Bayesian approach to single-cell differential expression analysis. Nat Methods 11, 740–742 (2014).

21. Budde, M. W. & Roth, M. B. The Response of *Caenorhabditis elegans* to Hydrogen Sulfide and Hydrogen Cyanide. Genetics 189, 521–532 (2011).

22. Vora, M. et al. The hypoxia response pathway promotes PEP carboxykinase and gluconeogenesis in *C. elegans*. Nat Commun 13, 6168 (2022).

23. Shen, C., Shao, Z. & Powell-Coffman, J. A. The *Caenorhabditis elegans rhy-1* gene inhibits HIF-1 hypoxia-inducible factor activity in a negative feedback loop that does not include vhl-1. Genetics 174, 1205–1214 (2006).

24. Pender, C. L. & Horvitz, H. R. Hypoxia-inducible factor cell non-autonomously regulates *C. elegans* stress responses and behavior via a nuclear receptor. eLife 7, e36828 (2018).

25. Satija, R., Farrell, J. A., Gennert, D., Schier, A. F. & Regev, A. Spatial reconstruction of single-cell gene expression data. Nat Biotechnol 33, 495–502 (2015).

26. Macosko, E. Z. et al. Highly Parallel Genome-wide Expression Profiling of Individual Cells Using Nanoliter Droplets. Cell 161, 1202–1214 (2015).

27. Hao, Y. et al. Integrated analysis of multimodal single-cell data. Cell 184, 3573–3587.e29 (2021).

28. Shen, C., Nettleton, D., Jiang, M., Kim, S. K. & Powell-Coffman, J. A. Roles of the HIF-1 Hypoxia-inducible Factor during Hypoxia Response in *Caenorhabditis elegans*. J. Biol. Chem. 280, 20580–20588 (2005).

29. Diehl, C. & Horvitz, H. R. The nonsense-mediated decay RNA-surveillance pathway facilitates the hypoxia response by C. elegans. Preprint at 10.64898/2026.05.06.721854 (2026).

30. ENCODE Project Consortium. An integrated encyclopedia of DNA elements in the human genome. Nature 489, 57–74 (2012).

31. Feng, D., Qu, L. & Powell-Coffman, J. A. Whole genome profiling of short-term hypoxia induced genes and identification of HIF-1 binding sites provide insights into HIF-1 function in *Caenorhabditis elegans*. PLoS One 19, e0295094 (2024).

32. Holdorf, A. D. et al. WormCat: An Online Tool for Annotation and Visualization of *Caenorhabditis elegans* Genome-Scale Data. Genetics 214, 279–294 (2020).

33. Feng, D., Qu, L. & Powell-Coffman, J. A. Transcriptome analyses describe the consequences of persistent HIF-1 over-activation in *Caenorhabditis elegans*. PLoS One 19, e0295093 (2024).

34. McGhee, J. D. The C. elegans intestine. in WormBook: The Online Review of C. elegans Biology [Internet] (WormBook, 2007).

35. Pukkila-Worley, R. & Ausubel, F. M. Immune defense mechanisms in the *Caenorhabditis elegans* intestinal epithelium. Curr Opin Immunol 24, 3–9 (2012).

36. Zimmer, M. et al. Neurons detect increases and decreases in oxygen levels using distinct guanylate cyclases. Neuron 61, 865–879 (2009).

37. Lewis, J. A., Wu, C.-H., Berg, H. & Levine, J. H. The Genetics of Levamisole Resistance in the Nematode *CAENORHABDITIS ELEGANS*. Genetics 95, 905–928 (1980).

38. Lewis, J. A., Wu, C. H., Levine, J. H. & Berg, H. Levamisole-resistant mutants of the nematode *Caenorhabditis elegans* appear to lack pharmacological acetylcholine receptors. Neuroscience 5, 967–989 (1980).

39. Fleming, J. T., et al. *Caenorhabditis elegans* Levamisole Resistance *Geneslev-1*, *unc-29*, and *unc-38* Encode Functional Nicotinic Acetylcholine Receptor Subunits. J Neurosci 17, 5843–5857 (1997).

40. Yuan, Y., Hilliard, G., Ferguson, T. & Millhorn, D. E. Cobalt Inhibits the Interaction between Hypoxia-inducible Factor-α and von Hippel-Lindau Protein by Direct Binding to Hypoxia-inducible Factor-α *. Journal of Biological Chemistry 278, 15911–15916 (2003).

41. Brenner, S. The genetics of *Caenorhabditis elegans*. Genetics 77, 71–94 (1974).

42. Mello, C. C., Kramer, J. M., Stinchcomb, D. & Ambros, V. Efficient gene transfer in *C.elegans*: extrachromosomal maintenance and integration of transforming sequences. EMBO J 10, 3959–3970 (1991).

43. Choi, H. M. T. et al. Third-generation in situ hybridization chain reaction: multiplexed, quantitative, sensitive, versatile, robust. Development 145, dev165753 (2018).

44. Rual, J.-F. et al. Toward improving *Caenorhabditis elegans* phenome mapping with an ORFeome-based RNAi library. Genome Res 14, 2162–2168 (2004).

45. Qadota, H. et al. Establishment of a tissue-specific RNAi system in *C. elegans*. Gene 400, 166–173 (2007).

46. Cao, J. et al. Comprehensive single cell transcriptional profiling of a multicellular organism. Science 357, 661– 667 (2017).

47. Altun, Z.F. et al. WormAtlas. *(ed.s) 2002-2024* https://www.wormatlas.org/hermaphrodite/muscleintro/MusIntroframeset.html.

48. Packer, J. S. et al. A lineage-resolved molecular atlas of *C. elegans* embryogenesis at single-cell resolution. Science 365, eaax1971 (2019).

49. Stuart, T. et al. Comprehensive Integration of Single-Cell Data. Cell 177, 1888–1902.e21 (2019).

50. Butler, A., Hoffman, P., Smibert, P., Papalexi, E. & Satija, R. Integrating single-cell transcriptomic data across different conditions, technologies, and species. Nat Biotechnol 36, 411–420 (2018).

51. Murray, J. I. et al. Multidimensional regulation of gene expression in the *C. elegans* embryo. Genome Res 22, 1282–1294 (2012).

52. Collins, K. M. & Koelle, M. R. Postsynaptic ERG Potassium Channels Limit Muscle Excitability to Allow Distinct Egg-Laying Behavior States in *Caenorhabditis elegans*. J Neurosci 33, 761–775 (2013).

